# Mutational impact of APOBEC3A and APOBEC3B in a human cell line and comparisons to breast cancer

**DOI:** 10.1101/2022.04.26.489523

**Authors:** Michael A. Carpenter, Nuri A. Temiz, Mahmoud A. Ibrahim, Matthew C. Jarvis, Margaret R. Brown, Prokopios P. Argyris, William L. Brown, Douglas Yee, Reuben S. Harris

**Author notes:** Correspondence to Reuben S. Harris. Equal contributions (Michael A. Carpenter, Nuri A. Temiz, Mahmoud A. Ibrahim).

## Abstract

A prominent source of mutation in cancer is single-stranded DNA cytosine deamination by cellular APOBEC3 enzymes, which results in signature C-to-T and C-to-G mutations in TCA and TCT motifs. Although multiple enzymes have been implicated, reports conflict and it is unclear which protein(s) are responsible. Here we report the development of a selectable system to quantify genome mutation and demonstrate its utility by comparing the mutagenic activities of three leading candidates - APOBEC3A, APOBEC3B, and APOBEC3H. The human cell line, HAP1, is engineered to express the *thymidine kinase* (*TK*) gene of HSV-1, which confers sensitivity to ganciclovir. Expression of APOBEC3A and APOBEC3B, but not catalytic mutant controls or APOBEC3H, triggers increased frequencies of *TK* mutation and nearly indistinguishable TC-biased cytosine mutation profiles in the selectable *TK* reporter gene. Whole genome sequences from *TK* mutant clones enabled an analysis of thousands of single base substitution mutations and extraction of local sequence preferences with APOBEC3A preferring YTCW motifs over 70% of the time and APOBEC3B just under 50% of the time (Y=C/T; W=A/T). Signature comparisons with breast tumor whole genome sequences indicate that most malignancies manifest intermediate percentages of APOBEC3 signature mutations in YTCW motifs, mostly between 50 and 70%, suggesting that both enzymes are contributing in a combinatorial manner to the overall mutation landscape. Although the vast majority of APOBEC3A- and APOBEC3B-induced single base substitution mutations occur outside of predicted chromosomal DNA hairpin structures, whole genome sequence analyses and supporting biochemical studies also indicate that both enzymes are capable of deaminating the single-stranded loop regions of DNA hairpins at elevated rates relative to control conditions. These studies combine to help resolve a long-standing etiologic debate on the source of APOBEC3 signature mutations in cancer and indicate that future diagnostic and therapeutic efforts should focus on both APOBEC3A and APOBEC3B.

## Introduction

Over the past decade, advances in DNA sequencing technologies and bioinformatics have helped to deconvolute a multitude of mutational processes that contribute to the genesis and evolution of cancer (see pan-cancer analysis by [1] and reviews by [2–4]). Through these approaches and complementary wet lab experiments, the APOBEC3 family of single-stranded (ss)DNA cytosine deaminases has emerged as one of the top three sources of single base substitution (SBS) mutation in cancer with particularly large contributions to tumors of the bladder, breast, cervix, lung, and head/neck. APOBEC3 signature mutations in cancer are defined as C-to-T transitions and C-to-G transversions in 5’-T<u>C</u>W motifs (W = A or T; SBS2 and SBS13, respectively) [1, 5–10]. This definition is conservative because several APOBEC3 enzymes can also accommodate 5’-<u>C</u>G and 5’-<u>methyl-C</u>G ssDNA substrates [9–17], which can also lead to C-to-T transition mutations and overlap with the mutation signature attributable to spontaneous water-mediated deamination of cytosine and methyl-cytosine nucleotides. Spontaneous, water mediated methyl-cytosine deamination is a clock-like mutation process that occurs predominantly in 5’-<u>methyl-C</u>G motifs and associates positively with a patient’s biological age (ageing signature [18]), whereas APOBEC3-catalyzed deamination is absent in most normal tissues, not associated with ageing, present in many primary tumors, and often enriched in metastases (APOBEC3 signature [1, 6, 8, 9, 18–23]).

The human APOBEC3 (A3) family is comprised of seven different enzymes with extensive homology and overlapping activities (reviewed by [24–26]), and it is unclear how much (or little) each contributes to the composite APOBEC3 mutation signature evident in tumor DNA sequences. The bulk of evidence favors two enzymes, APOBEC3A (A3A) and APOBEC3B (A3B), though to wildly different degrees depending on the study, and additional work has also implicated APOBEC3H (A3H). Evidence for A3A includes an intrinsic preference for 5’-TC substrates, high catalytic activity (highest of any human DNA deaminase), cell-wide localization, DNA damage responses, induction by the tumor virus HPV, and positive correlations between mRNA levels and tumor APOBEC3 mutation loads [11, 14, 16, 27–34]. Evidence for A3B includes all the same points except this enzyme is several-fold less active, localizes constitutively to the nuclear compartment, is induced by multiple DNA tumor viruses (herpesviruses, papillomaviruses, and polyomaviruses), and associates at both mRNA and protein levels with clinical outcomes [9, 11, 16, 19–21, 33–44]. Importantly, however, the APOBEC3 mutation signature can still accumulate in tumors that lack *A3B* due to a naturally occurring full gene deletion allele [45–47]. This observation with *A3B*-null tumors helped implicate a cell-wide, 5’-TC preferring variant of A3H (haplotype I) in contributing to the overall APOBEC3 mutation signature [46, 48, 49]. An important additional consideration is that A3A and A3B have been reported to exhibit broader tetranucleotide preferences upon expression in yeast, 5’-YTCW vs RTCW, respectively [50–52].

Here, we develop a human cellular system for mutation research and use it to compare the mutagenic potential of A3A, A3B, and A3H. The human cell line HAP1 was engineered to express a single copy of the HSV-1 *thymidine kinase* (*TK*) gene, which enables the drug ganciclovir to be used to select rare *TK* mutants and quantify mutation frequencies. Moreover, the *TK* gene can be amplified readily from ganciclovir-resistant (Gan^R^) clones by high-fidelity PCR and sequenced to provide initial assessments of mutation spectra (signatures) prior to undertaking additional experiments such as whole genome sequencing (WGS). Using this system, only expression of A3A and A3B cause significant increases in Gan^R^ mutation frequencies. Sanger sequences from panels of individual *TK* mutant clones show a clear TC-biased mutation pattern including two hotspots with no obvious hairpin structure. WGS of *TK* mutant clones demonstrates that both A3A and A3B can generate the APOBEC3 mutation signatures SBS2 and SBS13. These single base substitution mutations are mostly dispersed (non-clustered) throughout the genomes and both enzymes exhibit similar frequencies of mutation in TCW motifs in chromosomal DNA predicted to form non-hairpin versus hairpin structures. However, in comparison to catalytic mutant controls, both enzymes exhibit higher frequencies of APOBEC3 signature mutation in the single-stranded DNA loop regions of predicted hairpin structures. WGS also shows that A3A has a strong (slightly over 70%) preference for triggering cytosine mutations in YTCW sequence motifs, whereas A3B has a weaker (slightly under 50%) preference for the same motif. In comparison, APOBEC3 mutation signature-enriched primary breast cancers show predominantly intermediate frequencies of mutations in YTCW motifs, between 50% and 70%, suggesting involvement from both enzymes.

## Results

### HAP1-TK-M9 – a human cellular system to report DNA damage and mutagenesis

Model organisms such as *E. coli* and yeast are powerful systems for studying mutagens including DNA deaminases (*e.g*., original studies with A3 enzymes [34, 53–57]). However, these model organisms only recapitulate a subset of DNA repair and regulatory mechanisms found in human cells. We therefore sought to combine strengths of both approaches by introducing a single copy of a selectable reporter, the HSV-1 *thymidine kinase* (*TK*) gene, into the genome of the human cell line HAP1. Expression of the thymidine kinase (TK) protein confers exquisite sensitivity to the drug ganciclovir and, as for many antimicrobial agents, only *TK*-mutant, ganciclovir-resistant (Gan^R^) cells survive selection by this drug. Gan^R^ mutants can be characterized rapidly by conventional Sanger DNA sequencing because *TK* is a single open reading frame. Moreover, once informative Gan^R^ mutants are revealed by selection, secondary analyses including WGS can be used to uncover additional and potentially global features of a given mutation process.

The overall experimental workflow is shown in **Figure 1A**. To generate a “mother” clone of the commercially available HAP1 cell line, Sleeping Beauty (SB)-mediated transposition was used to introduce a single copy of a *TK-Neo* cassette into the genome [9]. Neo^R^ clones were selected with G418, expanded into healthy clonal populations (ca. 10^6^ cells/ml), and screened for ganciclovir sensitivity (Gan^S^). One mother clone, HAP1-TK-M9, was selected for further studies because it is Gan^S^, it is mostly diploid (apart from pre-existing chromosome aberrations), it cultures, engineers, and clones well (below), and it has a favorable *A3* expression profile (**Figure 1B**; **Figure 1 – Figure Supplement 1**). In particular, RT-qPCR measurements showed that its *A3A* and *A3B* mRNA levels are lower than those of the original parent line and that *A3H* mRNA levels are very low and near the detection threshold (**Figure 1B**). Genomic DNA sequencing also revealed that this cell line’s only *A3H* allele is haplotype III (ΔAsn15), which is known to produce an unstable protein [58–60].

**Figure 1.**
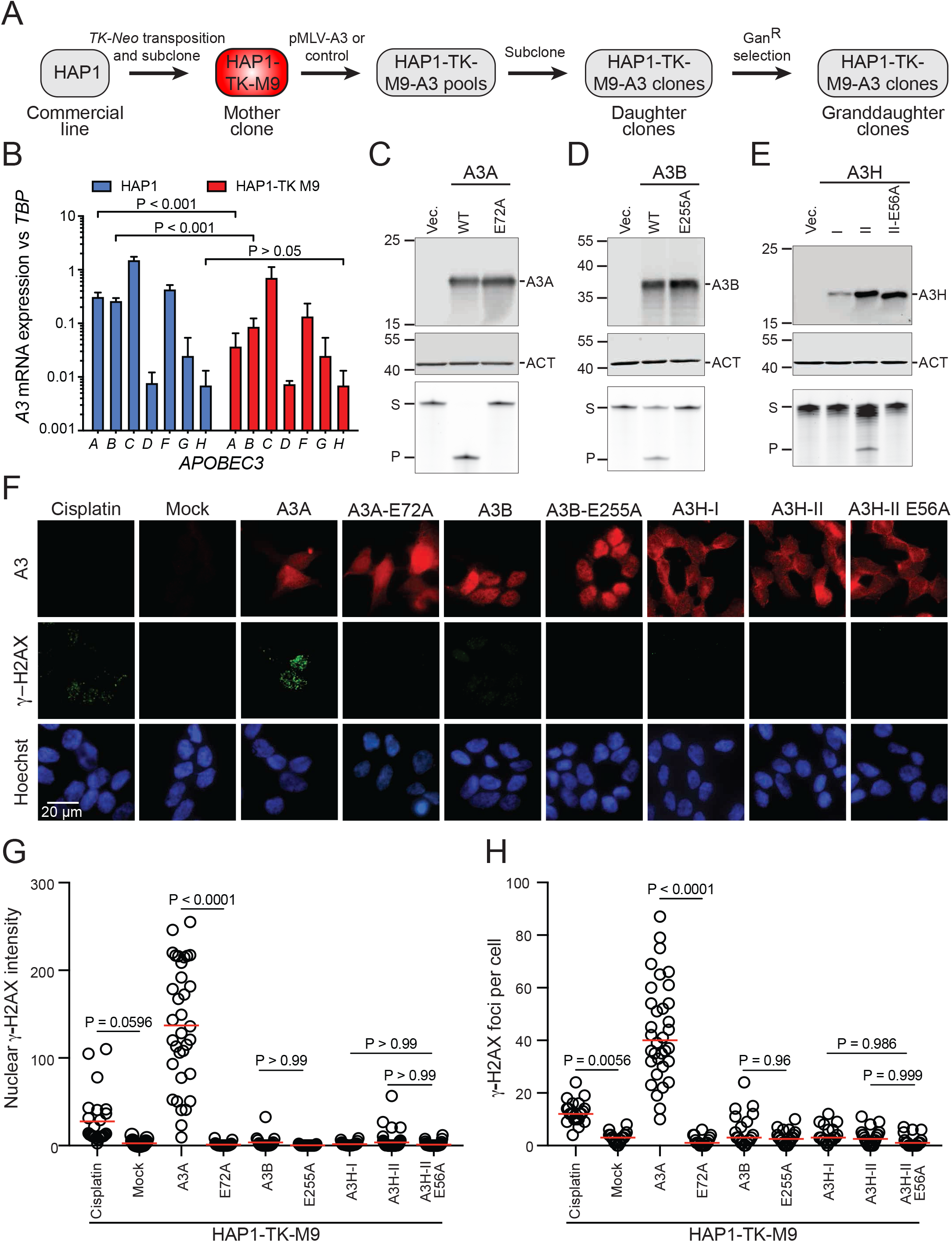
A3 activity in the HAP1-TK-M9 mutation reporter system. (**A**) Schematic of the construction of the HAP1-TK-M9 system and overall experimental workflow (see **Figure 1 – Figure Supplement 1** for HAP1-TK-M9 karyotype). (**B**) *A3* mRNA levels in parental HAP1 cells in comparison to the HAP1-TK-M9 daughter clone by RT-qPCR (p-values from Welch’s t-test). (**C-E**) Immunoblots of A3A, A3B, and A3H in WCE 24 hrs post-transfection of 293T cells. Primary antibodies are UMN-13, 5210-87-13, and P1-D8, respectively. Mouse anti-β-actin (ACT) is a loading control for the A3A and A3B blots, and rabbit anti-β-actin (ACT) for the A3H blot (see **Figure 1 – Figure Supplement 2** for anti-A3A UMN-13 mAb validation). The lower images show ssDNA deaminase activity of extracts from the same cell populations (S, substrate; P, product). (**F**) IF microscopy images of HAP1-TK-M9 cells expressing the indicated A3 enzymes or treated with 4 μM cisplatin for 24 hrs. A3 staining is red, γ-H2AX staining is green, and nuclei are blue from Hoechst (scale bar = 20 µm). (**G-H**) Quantification of pan-nuclear γ-H2AX intensity and discrete γ-H2AX foci, respectively, for each condition described in panel F (each data point represents an independent cell; n>25 cells per condition; p-values from one-way ANOVA test; red bar indicates mean for each group; representative of 2 biologically independent experiments).

### Functionality of human A3s expressed from MLV-based constructs

Plasmid constructs were assembled for these studies in which human *A3* expression is driven by a *MND* promoter from within an MLV-based retroviral construct. An additional feature of these constructs is a downstream puromycin resistance cassette, which facilitates selection of expressing cells. The functionality of each construct was assessed by transfection into 293T cells and, following 24-48 hrs incubation, protein analysis by immunoblotting and ssDNA deaminase activity assays (**Figure 1C-E**; custom rabbit anti-human A3A monoclonal antibody validation described in **Figure 1 – Figure Supplement 2**). Each A3 expressed at the expected kilodalton size and only the wildtype enzymes exhibited catalytic activity. A3H-I was expressed at lower levels than A3H-II and, accordingly, had less ssDNA deaminase activity in soluble extracts (**Figure 1E**), in agreement with prior reports [58–60].

We next used immunofluorescent (IF) microscopy to examine subcellular localization and DNA damage responses triggered by expression of each enzyme 3 days post-transduction into HAP1-TK-M9 cells (without puromycin selection). A3A appeared cell-wide and associated with a large increase in overall staining of the DNA damage marker γ-H2AX as well as an increase in individual γ-H2AX foci (representative images in **Figure 1F**, quantification in **Figure 1G-H**, and additional images in **Figure 1 – Figure Supplement 3**). Expression of an enzymatically inactive mutant, A3A-E72A, exhibited vector control levels of γ-H2AX staining. In comparison, neither expression of wildtype A3B nor A3H (haplotype I or II) was able to trigger statistically significant increases in γ-H2AX levels in HAP1-TK-M9 cells (representative images in **Figure 1F**, quantification in **Figure 1G-H**, and additional images in **Figure 1 – Figure Supplement 3**). These results were unexpected given that A3B localization is predominantly nuclear, and A3H (haplotypes I and II) can also access the nuclear compartment with some accumulation in nucleoli (**Figure 1F** and additional images in **Figure 1 – Figure Supplement 3**). These results differed from prior reports on A3B and A3H overexpression causing elevated γ-H2AX levels [9, 43, 48, 61], perhaps because expression levels here are lower with the *MND* promoter (versus strong Tet/Dox-inducible systems) and/or because the HAP1 system is somehow unique and more tolerant/adaptable to expression of these ssDNA deaminases. However, HAP1-TK-M9 cells are indeed capable of a canonical DNA damage response, as demonstrated by γ-H2AX accumulation following treatment with cisplatin (*cis*-diamminedichloroplatinum II; representative images in **Figure 1F**, quantification in **Figure 1G-H**, and additional images in **Figure 1 – Figure Supplement 3**).

To extend these results to a different cell line, our MLV-based A3 expression plasmids were transfected transiently into HeLa cells and, after 24 hrs incubation, subjected to additional analysis by IF microscopy. As with HAP1-TK-M9 cells, A3A appeared cell-wide (except nucleoli), A3B predominantly nuclear, and A3H haplotype II cytoplasmic with nucleolar accumulations (**Figure 1 – Figure Supplement 4A**). A3H haplotype I was not analyzed here due to low expression and weak ssDNA deaminase activity in HAP1-TK-M9 cells. Interestingly, A3A caused a strong pan-nuclear increase in γ-H2AX without concomitant focus formation (representative images in **Figure 1 – Figure Supplement 4A** and quantification in **Figure 1 – Figure Supplement 4B**). Similar results have been reported for A3A overexpression in other cell types [30, 62–64]. Interestingly, although a dose-responsive accumulation of γ-H2AX was expected given a wide range of transient transfection efficiencies for individual cells in each reaction, only a weak positive association was found with even low A3A staining cells exhibiting very high γ-H2AX levels (**Figure 1 – Figure Supplement 4B-C**). In contrast, comparatively low levels of nuclear γ-H2AX were observed in cells expressing A3B, A3H, and catalytic mutant derivative proteins, although A3B expression uniquely triggered a modest elevation of nuclear γ-H2AX levels in comparison to the background observed in cells expressing the corresponding catalytic mutant E255A protein (**Figure 1 – Figure Supplement 4A-D**). Moreover, as expected from the strong increase in γ-H2AX levels, only wildtype A3A caused significant increases in DNA breakage as quantified with alkaline comet assays (representative images in **Figure 1 – Figure Supplement 4E** and quantification in **Figure 1 – Figure Supplement 4F**). Taken together, results with two different cell lines demonstrate that A3A expression induces a strong DNA damage response (high γ-H2AX and DNA breakage), A3B expression triggers a modest DNA damage response (low γ-H2AX and no overt DNA breakage), and A3H or catalytic mutant A3A/B/H expression is indistinguishable from background levels in negative vector control conditions.

### *TK* mutation spectra of HAP1-TK-M9 with A3A, A3B, and A3H

To directly test which A3 enzymes cause genomic mutation in the HAP1-TK-M9 system, 24 independent single-cell derived daughter clones were obtained for A3A, A3B, A3H-I, and A3H-II expressing conditions as well as catalytic mutant and vector controls (**Methods**). A classical fluctuation analysis was performed by growing each single cell clone for 1 month to >10^7^ cells, subjecting each population to selection by ganciclovir, and allowing time for single Gan^R^ mutant cells to grow into countable colonies. Vector control conditions yielded a median Gan^R^ mutation frequency of 3 mutants per 5 million cells (mean = 3.6, SD = 2.8). In contrast, A3A- and A3B-expressing clones caused median Gan^R^ mutation frequencies to rise above 10 mutants per 5 million cells (A3A WT: median = 16, mean = 14, SEM = 2.2; A3B WT: median = 11, mean = 12, SEM = 1.8; **Figure 2A**). In comparison, expression of A3H-I or A3H-II or catalytic mutant derivatives of any of these DNA deaminases or empty vector controls failed to trigger increased Gan^R^ mutation frequencies (A3H-I WT: median = 3.0, mean = 3.6, SEM = 0.71; A3H-II WT: median = 2.0, mean = 2.8, SEM = 0.58; **Figure 2A**).

**Figure 2.**
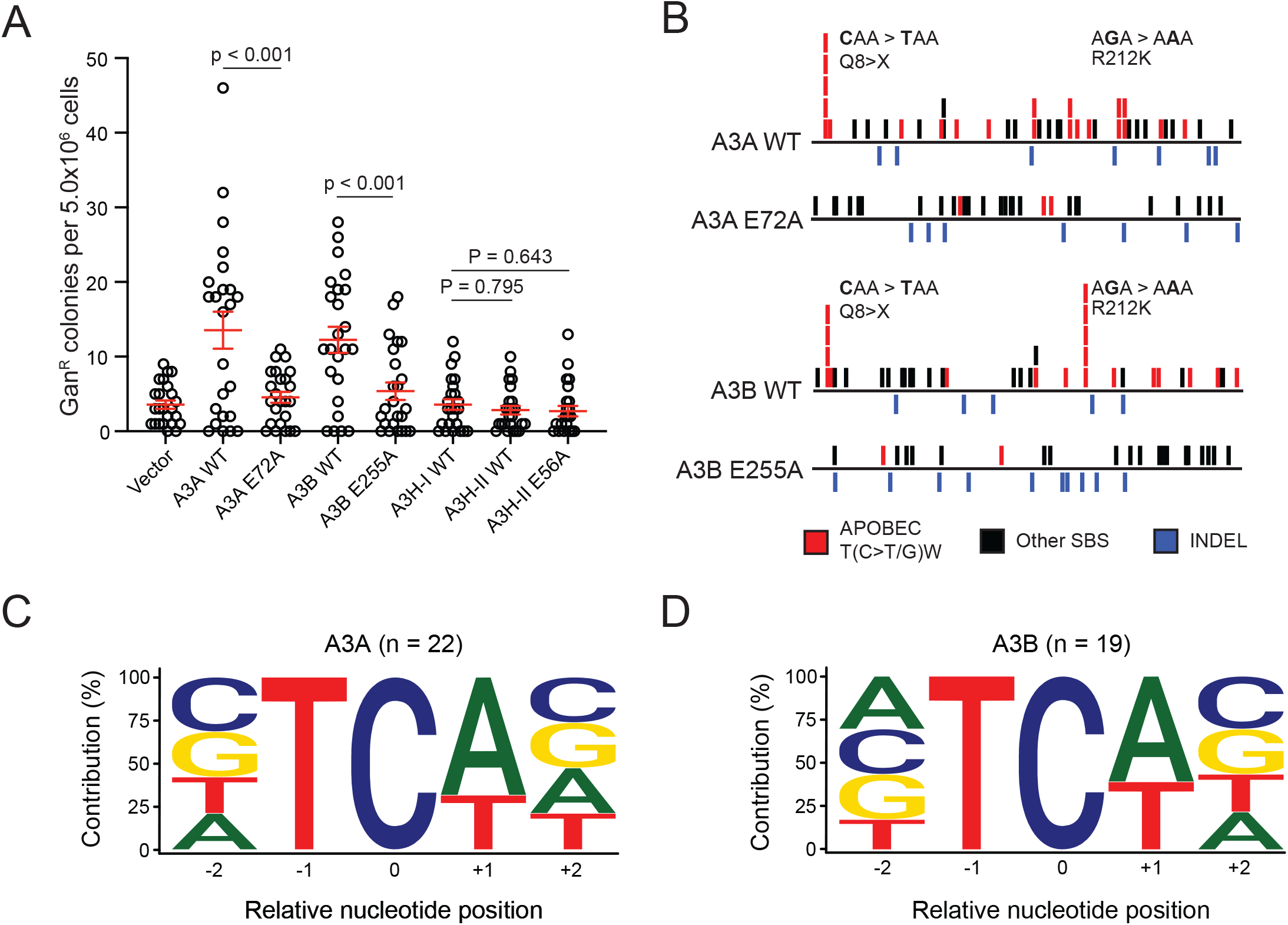
Characterization of *TK* mutations in ganciclovir-resistant clones. (**A**) A dot plot of Gan^R^ colonies generated under the indicated A3 expression or control conditions. Each data point represents the number of Gan^R^ mutants in a single clonal culture (mean +/- SD shown with p-values determined using Welch’s t-test). (**B**) Schematics representing all *TK* mutations observed under the indicated A3 expression conditions (APOBEC3 signature T(C>T/G)W mutations in red, other SBSs in black, and INDELs in blue; see **Figure 2 – Figure Supplement 1** for schematics of individual *TK* mutants for these and vector control conditions). Q8X and R212K mutation hotspots are labeled. (**C-D**) Pentanucleotide logos depicting the -2 and +2 sequence preferences flanking all T(C>T/G)W mutations that accumulated during A3A- or A3B-expression.

We next asked what types of genetic alterations led to inactivation of the *TK* gene in Gan^R^ granddaughter clones derived from the different A3-expressing and control conditions. The *TK* gene was PCR-amplified from genomic DNA of Gan^R^ granddaughter clones and Sanger sequenced. C-to-T and C-to-G mutations in an APOBEC3-signature trinucleotide motif (APOBEC), all other single base substitution mutations (other SBS), and all insertion/deletion mutations (INDELs) were placed into groups for comparison (red, black, and blue tics in **Figure 2B**, respectively; individual sequence schematics in **Figure 2 – Figure Supplement 1**). *TK* sequences derived from A3A-expressing granddaughter clones harbored a greater number of APOBEC3 signature mutations [12/20 clones contained at least 1 APOBEC mutation, 22 T(C>T/G)W mutations total, range of 0 to 3 SBS per sequence] relative to catalytic mutant control clones [3/19 clones contained 1 APOBEC mutation, 3 T(C>T/G)W mutations total, range of 0 to 1 SBS per sequence]. Similarly, *TK* sequences derived from A3B-expressing granddaughter clones also harbored a greater number of APOBEC3 signature mutations [11/20 clones contained at least 1 APOBEC mutation, 19 T(C>T/G)W mutations total, range of 0 to 3 SBS per sequence] relative to catalytic mutant control clones [2/18 clones contained 1 APOBEC mutation, 2 T(C>T/G)W mutations total, range of 0 to 1 SBS per sequence] (**Figure 2B**; **Figure 2 – Figure Supplement 1**). Additionally, *TK* cytosine bases 22 (Q8X) and 635 (R212K) emerged as candidate mutational hotspots in both the A3A- and A3B-expressing clones (**Figure 2B**; considered further in **Discussion**). No significant differences were found in the number of other SBS or INDEL mutations between A3A- or A3B-expressing clones and controls.

We next examined the broader sequence context of the 22 A3A- and 19 A3B-induced APOBEC3 signature mutations that occurred at 5’-TC dinucleotides in *TK* (**Figure 2C-D**). In both instances, A was preferred over T at the +1 nucleobase position relative to the mutated C, and this bias was not significantly different between the two enzymes (68% for A3A and 74% for A3B; p=0.367 by Fisher’s exact test). Similarly, no obvious biases were evident at the +2 or -2 nucleobase positions with all four nucleotides observed at similar frequencies for both enzymes. Moreover, even when pyrimidines and purines were grouped for comparison, A3A did not show an overt preference for C/T (Y) or A/G (R) at the -2 or +2 nucleobase positions (51% vs 49% and 52% vs 48%, respectively). Likewise, A3B also failed to show an overt preference for C/T (Y) or A/G (R) at the -2 or +2 nucleobase positions (45% vs 55% and 53% vs 47%, respectively). These similarities underscore the fact that small mutation numbers are primarily useful for delineating major signature differences such as the shifts described above from a heterogeneous pattern in catalytic mutant- or vector control-expressing cells towards a predominantly 5’-TC focused SBS mutation pattern in A3A- and A3B-expressing cells.

### WGS shows SBS2 and SBS13 signatures reflecting the intrinsic biochemical preferences of A3A and A3B

To prepare for WGS of Gan^R^ granddaughter clones, RNA sequencing was done to confirm expression of each exogenously expressed *A3* construct and compare mRNA levels relative to established breast cell lines and primary breast tumors. All expression values were determined relative to those of the conserved housekeeping gene *TBP* to be able to compare RNAseq data from different sources (*e.g.*, unrelated cell lines and tumors). Interestingly, the average *A3A* and *A3A-E72A* mRNA levels in Gan^R^ granddaughter clones was over 5-fold higher than the average endogenous *A3A* expression levels of APOBEC3 signature-enriched breast cancer cell lines BT474 and MDA-MD-453, breast cancer cell lines of the CCLE, or breast tumors of TCGA (**Figure 3 – Figure Supplement 1A**). It is possible that A3A levels are higher in this system due to difficulty identifying a sufficiently weak promoter for constitutive expression. In comparison, *A3B* and *A3B-E255A* mRNA levels in Gan^R^ granddaughter clones are similar to the averages reported for breast cancer cell lines of the CCLE and breast tumors of the TCGA, and approximately 2-fold lower than those of BT474 and MDA-MD-453 (**Figure 3 – Figure Supplement 1A**). *A3H* (haplotype I) mRNA levels showed greater variance but only two clones were analyzed by RNAseq and WGS due to negative results above with IF microscopy experiments and *TK* mutation analysis.

**Figure 3.**
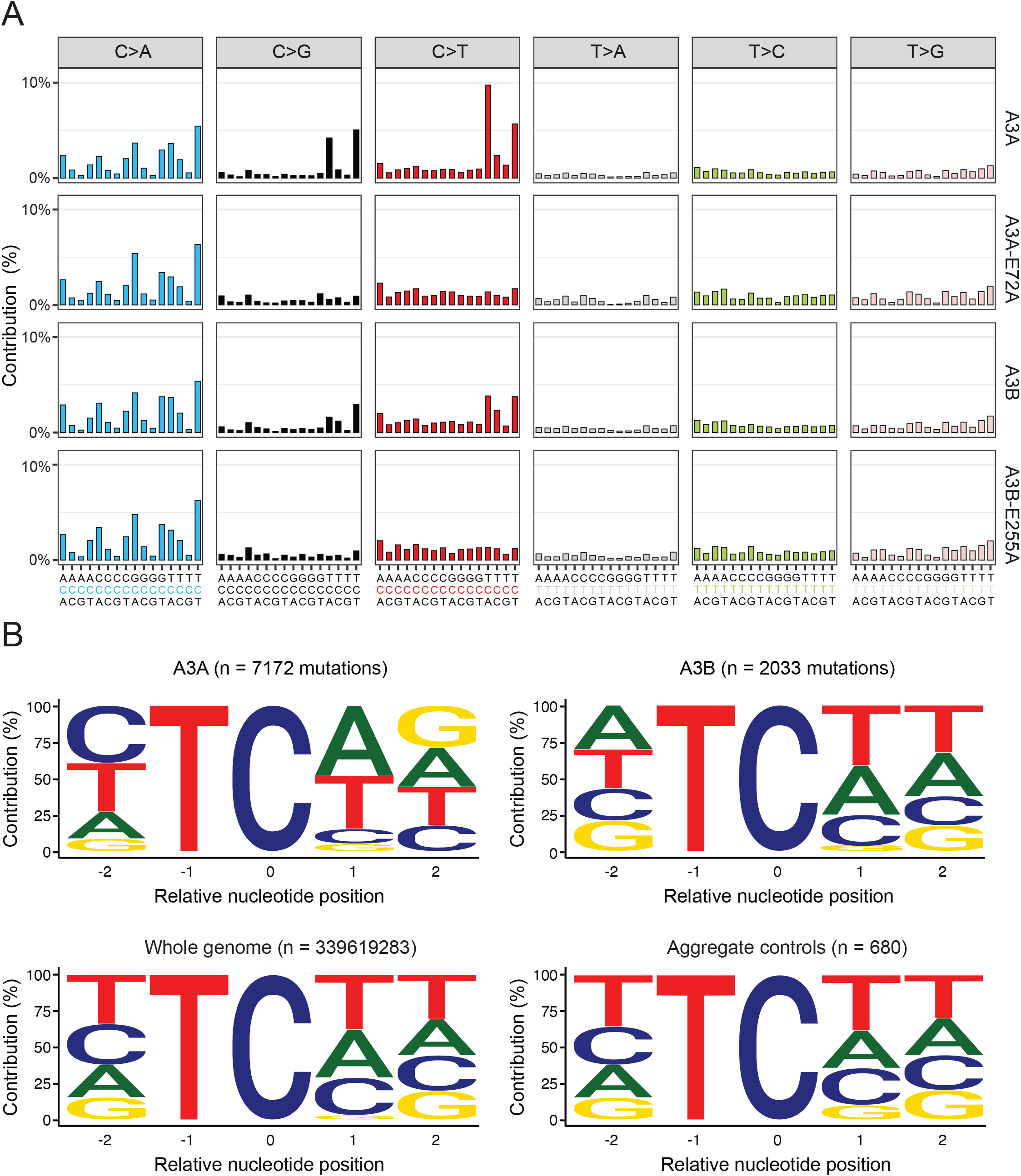
Single base substitution mutation signatures in genomes of ganciclovir-resistant clones. (**A**) Trinucleotide profiles of pooled SBSs across all clones sequenced for each listed experimental condition (A3A n=6; A3A-E72A n=2; A3B n=5; A3B-E255A n=2). See **Figure 3 – Figure Supplement 1 and 2** for mRNA and protein level expression confirmation, respectively, and **Figure 3 – Figure Supplement 5** for SBS profiles from each WGS. (**B**) Pentanucleotide logos depicting -2, +1 and +2 sequence preferences flanking all C-to-T and C-to-G mutated TC motifs in WGS from HAP1-TK-M9 cells expressing A3A or A3B in comparison to aggregate controls (catalytic mutants and GFP only conditions, which do not show evidence for APOBEC3 signature mutations; **Figure 3 – Figure Supplement 5**). The distribution of nucleobases flanking all TC motifs in the human genome is shown for comparison.

The mRNA expression levels of the other four *A3* genes, as well as *AICDA* (*AID*), *APOBEC1*, *APOBEC2*, and *APOBEC4*, were also quantified and compared with those of *A3A*, *A3B*, and *A3H* (**Figure 3 – Figure Supplement 1B**). Endogenous *A3C* was expressed at similarly high levels in all granddaughter clones, providing a stable relatively internal control. Endogenous *A3F* and *A3G* were expressed at lower but still detectable levels, and endogenous *AICDA*, *APOBEC1*, *APOBEC2*, *APOBEC4*, and *A3D* were expressed at very low or undetectable levels. As expected, levels of ectopically expressed *A3A*, *A3B*, and *A3H* mRNA exceeded those in the HAP1-TK-M9 parent clone as well as those in vector expressing granddaughter controls. In addition, protein expression of A3A, A3B, and A3H was confirmed in *TK* mutant granddaughter clones by immunoblotting and activity by ssDNA deamination assays (**Figure 3 – Figure Supplement 2**). Finally, the integration sites of the *TK* reporter and MLV-based A3 expression constructs were determined using WGS reads. Data from multiple clones demonstrated a single *TK-Neo* integration site in chromosome 3 between nucleotides 143996100-143996500, which is ∼3.5 kbp downstream of the nearest gene (*DIPK2A*), and representative *A3* expressing clones additionally have single MLV-A3 insertion sites on different chromosomes as expected from the independent reactions and low multiplicities of infection that used initially to establish these clones (**Figure 3 – Figure Supplement 3**).

To investigate mutational differences genome-wide, Illumina short-read WGS was done for randomly selected Gan^R^ granddaughter clones (specified by “WGS” in **Figure 2 – Figure Supplement 1**). Mutations unique to each granddaughter were identified by calling SBS variations versus the genomic DNA sequence of the HAP1-TK-M9 mother clone. This approach eliminated any somatic variation that accumulated in the Gan^S^ mother clone prior to transduction with each A3 or control expression construct. Thus, all new SBS mutations had to be present in a significant proportion of reads from the Gan^R^ granddaughter clones and absent from the reads from the Gan^S^ mother clone and, as such, must have occurred in the presence of an active A3 enzyme or a catalytic mutant control.

In A3A-expressing Gan^R^ granddaughter clones, the total number of unique SBSs ranged from 2057 to 5256 (n=6, median=3129, mean=3494, SD=1358; **Figure 3 – Figure Supplement 4A**). The total number of SBSs in A3B-expressing clones was lower, ranging from 1920 to 2652 (n=5, median=2346, mean=2334, SD=278). In comparison, the total number of SBSs in catalytic mutant and GFP control Gan^R^ granddaughter clones ranged from 1230 to 2204 (n=6; median=1748, mean=1729, SD=388). SBS mutations are further broken down into those occurring within NCN and TCW motifs in dot plots in **Figure 3 – Figure Supplement 4B-E**. Most importantly, analyses of the trinucleotide contexts of all unique SBS mutations revealed strong C-to-T and C-to-G mutation biases in 5’-TCA and 5’-TCT motifs in A3A-expressing clones and weaker, but still significant, mutation biases in the same motifs in A3B-expressing clones (**Figure 3A**; **Figure 3 – Figure Supplement 4C and E**; **Figure 3 – Figure Supplement 5**). In other words, only A3A- and A3B-expressing granddaughter clones exhibited significant amounts of APOBEC3 signature single base substitution mutations.

This result was confirmed by assessing APOBEC3 signature enrichment scores [50, 65], which indicated that 6/6 A3A-expressing clones and 4/5 A3B-expressing clones have significant enrichments of APOBEC3 signature mutations, whereas clones expressing catalytically inactive A3A or A3B, as well as clones expressing A3H-I or vector control have none (**Figure 3 – Figure Supplement 6A**). A complementary bioinformatics analysis, non-negative matrix factorization (NMF [66]), yielded similar results with “signature A” resembling SBS2 and SBS13 in A3A- and A3B-expressing clones (*i.e*., APOBEC3 signature) and “signature B” occurring in all clones regardless of A3 presence or functionality (**Figure 3 – Figure Supplement 6B**). In comparison, genome-wide patterns of insertion/deletion mutations (INDELs) appeared largely unaffected by A3A or A3B (**Figure 3 – Figure Supplement 7**), in agreement with aforementioned *TK* mutation data where the majority of Gan^R^ mutations are single base substitution mutations and the INDEL frequencies were similar for A3A, A3B, and catalytic mutant conditions.

We next analyzed the broader contexts of the 5’-TC-focused C-to-T and C-to-G single base substitution mutations that accumulated in A3A- and A3B-expressing clones (n=7172 and n=2033, respectively) in comparison to those that accumulated in aggregate control clones (n=680) as well as the overall distribution of 5’-TC in the human genome (n=339619283) (**Figure 3B** and **Supplementary Table S1**). First, a bias for +1 A over +1 T emerged in A3A-expressing clones (48.3% > 36.3%), whereas the opposite bias was evident in A3B-expressing clones (33.9% < 41.2%). However, for both enzymes, the percentage of +1 A and T (W) was similar (84.6% and 75.1%, respectively). Second, no significant bias was noted at the +2 position except that guanine is slightly over-represented in pentanucleotide motifs derived from A3A-expressing conditions compared to motifs derived from A3B-expressing clones. Third and most importantly, an exceptionally strong bias for a pyrimidine nucleobase (C or T) occurred at the -2 position in A3A-expressing clones (72.9% YTC/NTC), which was also evident in the broader APOBEC3 tetranucleotide context (72.7% YTCW/NTCW). This cytosine mutation preference resembles the strong -2 pyrimidine bias reported for human A3A in murine hepatocellular carcinomas [67], the chicken B cell line DT40 [73], and yeast [50, 51]. Fourth, in contrast, a slight bias for purine nucleobases (A or G) was apparent at the -2 position in A3B-expressing clones (51.1% RTC/NTC), which was also reflected in the broader APOBEC3 tetranucleotide context (52.8% RTCW/NTCW). This latter result also agreed with prior data from yeast [50, 51], but contrasted slightly with the mutation signature detected in tumors derived from human A3B-expresssing mice (47% RTCW [68]). Possible explanations for this variability are considered in **Discussion**.

### Features of A3A and A3B mutagenesis in the HAP1-TK-M9 system

X-ray structures have revealed a U-shaped bend in ssDNA substrates bound by A3A and A3B [16, 29], and other studies have indicated that similarly bent ssDNA loop regions of hairpins (*i.e*., DNA cruciform or stem-loop structures) may be preferred substrates for deamination by A3A [52, 69, 70]. To ask whether this preference extends to the HAP1-TK-M9 system described here, we analyzed our A3A and A3B *TK* PCR sequences and granddaughter clone WGS data for evidence of mutagenesis in the single-stranded loop regions of DNA hairpin structures. First, neither of the two A3A/B mutation hotspots in the *TK* gene reported above appeared to be part of predicted stem-loop structures. Second, none of the top-100 cruciform structures reported previously to harbor recurring APOBEC3 signature mutations in tumors [52] were mutated in our HAP1-TK-M9 WGS data sets. Third, in global comparisons of base substitution mutations, the frequency of APOBEC3 signature TCW mutations was similar in ssDNA loop regions of predicted hairpin structures versus non-hairpin regions (*i.e*., APOBEC3 signature mutation events were not enriched in the loop regions of stem-loop structures over those occurring in canonical ssDNA substrates; **Figure 4 – Figure Supplement 1**). Moreover, the frequency of APOBEC3 signature mutations in hairpin or non-hairpin structures appeared similar (not statistically distinguishable) in A3A- and A3B-expressing conditions. However, interestingly, the frequency of APOBEC3 signature mutations in predicted loop regions of hairpin structures appeared higher in A3A and A3B expressing clones in comparison to all non-catalytic control conditions (**Figure 4 – Figure Supplement 1**; P = 0.0655 and P = 0.0367 by Welch’s t-test, respectively). The A3A versus control comparison likely failed to reach statistical significance due to the small number of clones with WGS (n = 6) and the large variance in numbers of TCW mutations in hairpin loop regions in the different clones (range = 1-16 TCW mutations). Nevertheless, these results combined to indicate that single-stranded loop regions of hairpin structures in human chromosomal DNA may be similarly susceptible to deamination by both A3A and A3B.

To assess relative rates of A3A and A3B-catalyzed deamination of experimental hairpin versus non-hairpin substrates, we used purified enzymes and ssDNA substrates representing two previously reported RNA editing hotspots of A3A – *SDHB* and *NUP93* [52, 71]. The *SDHB* hairpin is predicted to have a 5 bp stem and a 4 nt loop, and the *NUP93* hairpin a 7 bp stem and a 4 nt loop (**Figure 4A-B**). The control oligonucleotides have the same loop region sequences and a randomization of one-half of the nucleobases in the hairpin stem to reduce base-pairing potential. In each case, the linear substrates migrated similarly on native and denaturing PAGE while the hairpin substrates migrated faster by native PAGE and similar to the linear substrates when denatured, thereby confirming the integrity of both hairpins (**Figure 4 – Figure Supplement 2**). A3A and A3B were affinity-purified from human cells and incubated under single hit conditions with these oligonucleotide substrates over time. First, both A3A and A3B showed a strong preference for deaminating the *SDHB* hairpin substrate in comparison to a linear control with the same nucleobase content scrambled (∼4- and ∼8-fold preference, respectively; **Figure 4A**). A3A showed higher rates of deamination than A3B on both the hairpin and the linear substrate in agreement with prior studies [52]. Thus, the relative deamination rates for *SDHB* substrates were: A3A/hairpin > A3B/hairpin > A3A/linear > A3B linear (120, 41, 31, and 5.1 nM/min, respectively). However, a different picture emerged from analyses of deamination of *NUP93*-based substrates **Figure 4B**). Rates of A3A-catalyzed deamination were similarly high for the *NUP93* hairpin and linear control with the same nucleobase content scrambled. In contrast, A3B showed higher rates of deamination of the linear substrate and was only able to deaminate the hairpin substrate with low efficiencies and linear kinetics. Thus, the relative deamination rates for *NUP93* substrates are: A3A/hairpin = A3A/linear >> A3B linear > A3B/hairpin (93, 83, 7.7, and 1.3 nM/min, respectively). These results combined to show that *<u>both</u>* A3A and A3B can deaminate hairpin and linear ssDNA substrates and, further, that it is possible to identify sites such as the *NUP93* hairpin (here with DNA and previously shown with RNA and DNA [52]) that are strongly (and perhaps even exclusively, in a few instances) preferred by a single A3 enzyme.

**Figure 4.**
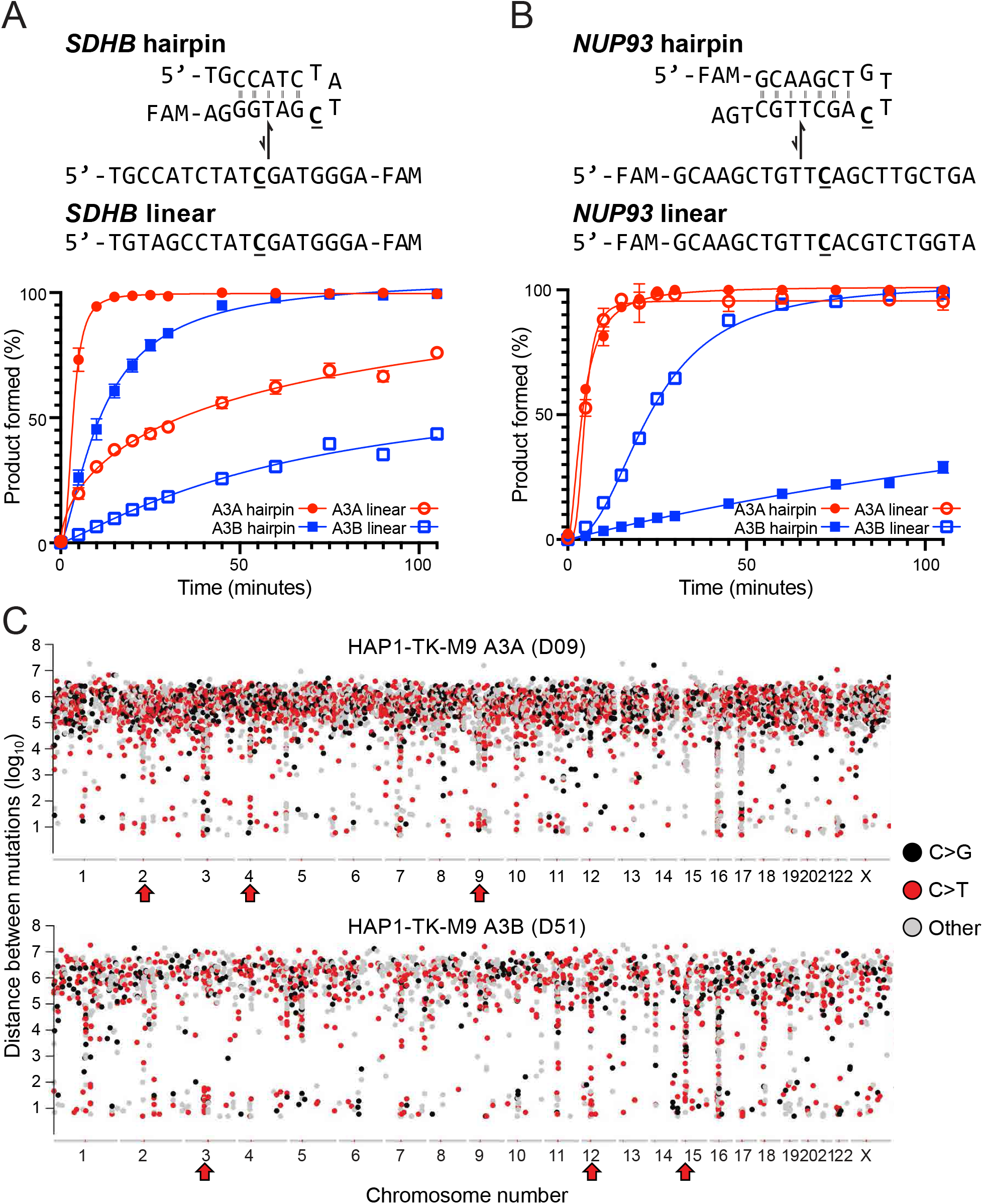
Mesoscale properties of A3A and A3B *in vitro* and in the HAP1-TK-M9 system. (**A-B**) Deamination kinetics of A3A and A3B using *SDHB* and *NUP93* DNA hairpin substrates in comparison to corresponding linear controls made by scrambling the 5’ or 3’ portion of the stem, respectively. See text for full description and **Figure 4 – Figure Supplement 1** for a genome-wide analysis and **Figure 4 – Figure Supplement 2** for a gel-based confirmation of DNA oligonucleotide integrity. (**C**) Rainfall plots of genome-wide intermutation distances showing APOBEC3 signature kataegic tracts (red arrows) in representative A3A- and A3B-expressing Gan^R^ clones (C>T mutations are red, C>G black, other SBS gray). Kataegis are defined here as ≥2 strand-coordinated APOBEC3 signature SBS mutations within a 10 kbp window.

Another feature of APOBEC3 mutagenesis in human cancer is clusters of strand-coordinated cytosine SBS mutations in TCA and TCT motifs most likely caused by processive deamination of exposed tracts of ssDNA (aka. *kataegis*; here defined as ≥2 strand-coordinated APOBEC3 signature mutations within a 10 kbp window) [5, 7, 66, 72]. No APOBEC3 signature *kataegic* events were observed in control conditions. However, several APOBEC3 signature *kataegic* events were evident in the genomic DNA of both A3A and A3B expressing granddaughter clones (*e.g*., **Figure 4C**). For instance, one A3A-attributable *kataegic* event was comprised of 7 T(C>T/G)W mutations, and an A3B-attributable *kataegic* event included 8 T(C>T/G)W mutations. Interestingly, however, the frequency of *kataegic* events did not differ significantly between A3A- and A3B-expressing granddaughter clones (A3A: median = 30, mean = 35 events, SD = 8.3; A3B: median = 12, mean = 15, SD = 6.9; p = 0.28 by Welch’s t-test). These results show that both A3A and A3B can cause *kataegis* in a human cell line and, surprisingly, at similar overall frequencies.

### APOBEC signature etiology in primary breast tumors

Sequencing data from model systems such as HAP1-TK-M9 are powerful because the resulting mutation signatures can help to establish cause-and-effect relationships for comparison to more complex tumor WGS data sets to identify similarities and, potentially, to infer the precise source of an observed mutation signature in individual tumors. We therefore performed an unsupervised clustering analysis to compare the pentanucleotide cytosine mutation signatures derived from sequencing A3A, A3B, and A3H expressing HAP1 clones and those from primary breast tumors with WGS available through the ICGC data portal resource. This analysis revealed three distinct tumor groups with respect to APOBEC3 signature mutations: 1) a group that showed similarity to A3A-expressing HAP1-TK-M9 clones, 2) a group that showed similarity to A3B-expressing HAP1-TK-M9 clones, and 3) a group that showed little to no significant APOBEC3 mutation signature (**Figure 5**). As expected from a lack of substantial APOBEC3 signature mutations above in the *TK* reporter or in representative WGS, A3H-I expressing clones and catalytic-inactive clones clustered with non-APOBEC3 signature tumors.

**Figure 5.**
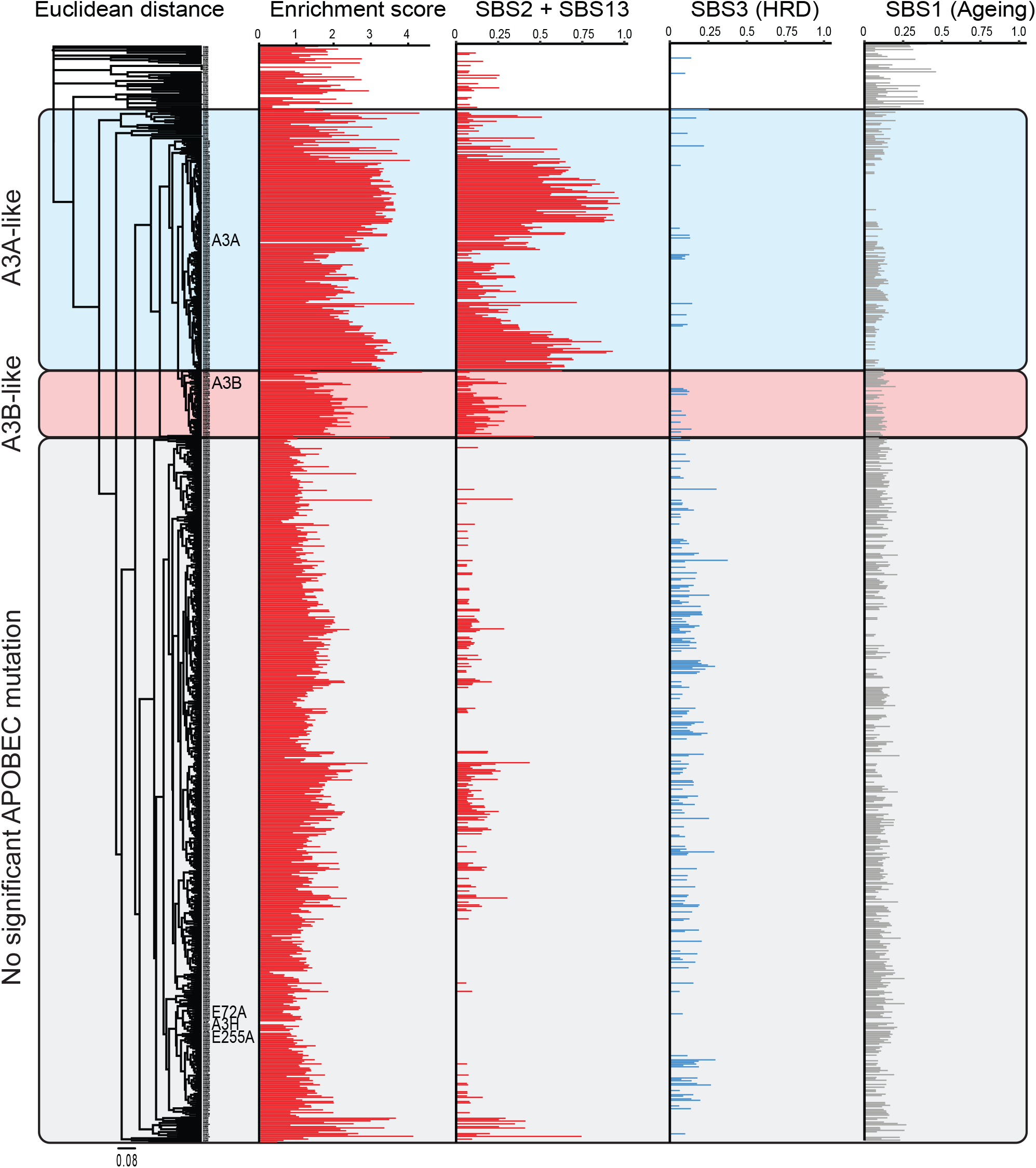
A composite origin of APOBEC3 signature mutations in breast cancer. An unsupervised clustering analysis of similarity between the pentanucleotide SBS profiles from WGSs of the A3A, A3B, and control Gan^R^ clones described here versus those from primary breast tumor whole-genome sequencing data sets (ICGC, n=784). The APOBEC3 mutation signature is represented by both enrichment score and SBS2+13 (red), HRD signature as SBS3 (blue), and ageing signature as SBS1 (gray).

Both the A3A-like and A3B-like groups were comprised of tumors that show significant levels of APOBEC3 signature mutations and correspondingly high enrichment scores (**Figure 5** and **Supplementary Table S1**). However, only a small proportion of the A3A-like group of breast tumors showed ≥71.7% APOBEC3 signature mutations in YTCW motifs, which is the overall preference of A3A observed here in the HAP1-TK-M9 system. Similarly strong YTCW preferences have also been reported recently for A3A expression in the chicken B cell line DT40 [73] and for A3A-driven murine hepatocellular carcinomas [67]. These observations therefore suggest that A3A alone may not account for the observed composite APOBEC3 mutation signature in individual breast tumors and, moreover, that many tumors in A3A-like group may include contributions from another A3 enzyme, most likely A3B. As regards the A3B-like group, most tumors have higher percentages of C-to-T and C-to-G mutations in YTCW motifs than can be explained by A3B alone. For instance, most tumors manifest a greater proportion of YTCW mutations than observed here for A3B in the HAP1-TK-M9 system (47.2%) and for A3B-driven tumors in mice (53%) [68]. These intrinsic preferences therefore combined to suggest that the observed percentages of APOBEC3 signature mutations in most breast tumors may be a composite resulting from the combinatorial activities of both A3A and A3B activity. In comparison, the homologous recombination repair deficiency (HRD) signature (SBS3) appeared underrepresented in both the A3A- and A3B-like tumor groups, and the ageing signature (SBS1) occurred in all three groups regardless of the presence or absence of an APOBEC3 mutation signature enrichment (**Figure 5**).

## Discussion

Here we have reported the development and implementation of a genetic system to investigate mutational processes in the human HAP1 cell line. Like many bacterial and yeast model systems, the HAP1-TK-M9 system enabled a uniform cytotoxic selection with ganciclovir such that only *TK* mutant cells survive. An analysis of clonally derived, A3A and A3B expressing *TK* mutants by Sanger sequencing of high-fidelity PCR amplicons demonstrated a strong shift in the mutational pattern from a variety of different base substitution mutations in control conditions to a strongly 5’TCW-biased pattern (**Figure 2**). Interestingly, the *TK* mutation spectra inflicted by A3A and A3B were very similar including two shared hotspots (Q8X, R212K) and no obvious mesoscale features such as palindromic sequences capable of hairpin formation. This result may be due to the limited number of mutable cytosines and TC motifs in *TK* that confer resistance to ganciclovir (local base composition) and/or to selective pressure. Regardless of the precise molecular explanation, an analogy can be drawn with the mutational spectrum of the *PIK3CA* gene in breast, head/neck, cervical, and others cancers, which has two prominent APOBEC3 mutation hotspots (E542K, E545K) and no obvious hairpin structures [52, 74]. These observations combined to suggest that selective pressure has the potential to overshadow the intrinsic preferences of individual APOBEC3 enzymes and complicate assignment of direct cause-and-effect relationships.

Drawing direct connections between A3A and/or A3B and a given mutation, even a prominent hotspot, has been additionally challenging due to the fact that both enzymes can deaminate DNA cytosines in linear substrates as well as single-stranded loop regions of stem-loop structures and, importantly, rates can vary dramatically between different substrates (*e.g*., **Figure 4** and prior biochemical studies [52, 69, 70]). Thus, the *TK*-based system described here is capable of yielding informative, rapid, and inexpensive mutation data sets with positive results motivating genome-wide analyses where mutations are unselected and larger mutation data sets enable broader analyses. For instance, our unbiased WGS analysis of A3A- and A3B-expressing clones indicated that both enzymes are capable of deaminating the ssDNA loop regions of hairpin substrates at similar frequencies (**Figure 4 – Figure Supplement 1**). This result was somewhat anticipated by structural studies where both A3A and a mutant A3B catalytic domain were shown to bind to ssDNA in a U-shaped conformation [16, 29], but it was also unexpected due to recent reports describing strong RNA and DNA hairpin biases for A3A [52, 69]. Additional biochemical and WGS studies will be required to confirm these results as well as extend them to additional cell lines and experimental systems. One potential drawback of the HAP1-TK-M9 system is that an exclusive focus on Gan^R^ clones might overestimate the mutational impact of a given process. However, earlier work also used HAP1 cells to successfully characterize a variety of different mutation sources, but these studies did leverage the power of a lethal genetic selection nor did they address the enzyme(s) responsible for APOBEC3 signature mutations [75, 76].

Taken together, the HAP1-TK-M9 studies here have demonstrated unambiguously that both A3A and A3B can inflict an APOBEC3 mutation signature in human genomic DNA with, in both instances, ssDNA deamination events immortalizing predominantly as C-to-T and C-to-G mutations in TCA and TCT trinucleotide motifs (**Figure 2**, **Figure 3**, and **Supplementary Table S1**). Over identical month-long timeframes, A3A causes 4-fold more APOBEC3 signature mutations in comparison to A3B (6070 vs 1528 mutations from 6 and 5 subclone WGSs, respectively). This difference may be explained in part by super-physiological A3A expression levels in the HAP1-TK-M9 system (at least 5-fold higher than levels in breast tumors or cell lines) and in part by the higher intrinsic activity of this enzyme in comparison to A3B. Regardless, both A3A and A3B inflicted thousands of TC-focused APOBEC3 signature mutations, which enabled comparisons between extended intrinsic preferences. Most importantly, A3A has a strong preference for a pyrimidine at the -2 position relative to the target cytosine (72.7% YTCW). This - 2 pyrimidine bias mirrors original results from human A3A expression in yeast [50, 51], as well as recent WGS results from human A3A expression in the chicken B cell line DT40 (∼70% YTCW [73]) and from human A3A-induced murine hepatocellular carcinomas (70% YTCW [67]). Also, similar to original studies in yeast [50, 51], human A3B showed a slight negative preference for YTCW motifs (47.2% YTCW) and a corresponding enrichment for RTCW motifs (52.8% RTCW; **Figure 3B**, **Figure 3 – Figure Supplement 4**, and **Supplementary Table S1**). In comparison, recent studies showed that human A3B driven tumors in mice (hepatocellular carcinomas and B cell lymphomas) exhibit an opposite APOBEC3 mutation signature bias with 53% YTCW and 47% RTCW [68]. These differences in APOBEC3B local preferences in the different systems may be due to genetic and/or epigenetic factors including but not limited to different base content, chromatin states, DNA repair processes, and/or post-translational regulatory mechanisms. They could also be due to simple stochastic variation and, accordingly, we hypothesize that A3B is non-discriminatory at the -2 position relative to the target cytosine and that this may relate to the amino acid composition of catalytic domain loop 1 residues relative to those of A3A (loop 3, loop 7, and most other active site residues are identical or nearly identical).

Regardless of the precise molecular explanation(s) for differences between experimental systems, the fact that both A3A and A3B can inflict YTCW mutations in human cells helps to inform interpretations of the APOBEC3 enzyme responsible for the overall APOBEC3 mutation program in cancer. For instance, based on comparisons of A3A- and A3B-attributable single base substitution mutation signatures observed here in the HAP1-TK-M9 system and extracted APOBEC3 mutation signatures from 784 breast cancer WGS, it is likely that neither enzyme’s preferred motif fully explains the composite signature in most individual tumors (**Figure 5** and **Supplementary Table S1**). As regards A3A, the majority of individual breast tumors has ≤72.7% APOBEC3 signature mutations in YTCW motifs, which is the overall preference of A3A here in the HAP1-TK-M9 system suggesting the involvement of at least one other A3 enzyme. As regards A3B, most individual breast tumors have a larger proportion of mutations in YTCW motifs than can be explained by A3B alone (≥47.2%). Thus, it is likely that both A3A and A3B contribute to the composite single base substitution mutation signature observed in individual APOBEC3 signature enriched breast tumors. However, we cannot exclude the possibility that some breast tumors may be mutated exclusively by A3A or A3B due to factors listed above, or that some breast tumors may have small mutagenic contributions from other APOBEC3 enzymes. Of course, A3B is not a direct factor in *A3B*-null breast tumors, although the inherited deletion that removes all *A3B* coding sequences may dysregulate *A3A* expression [45, 46]. In addition, the WGS data sets for A3A- and A3B-expressing HAP1-TK-M9 clones described here may be useful for comparing with APOBEC3 attributable events in other tumor types.

In addition to selective pressures and mesoscale features, additional factors are likely to influence the APOBEC3-attributable fraction of an overall tumor mutational landscape including whether A3A and/or A3B is expressed, expression levels, duration of expression, intrinsic activity, and accessibility of chromosomal DNA (replication stress, R-loop levels, chromatin state, *etc*.). With regards to studies here with the HAP1-TK-M9 system, both A3A and A3B were expressed constitutively from the same promoter/construct for identical durations prior to ganciclovir selection, A3A is intrinsically more active than A3B, A3A is cell-wide and A3B predominantly nuclear, and yet these enzymes and cellular factors combined to yield remarkably similar *TK* mutation frequencies and only a 4-fold difference in overall genome-wide SBS mutation level. With respect to cancer, the *A3A* gene is expressed at lower levels than *A3B* in almost all cell lines and tumors, A3A is cell-wide or predominantly cytoplasmic where A3B is constitutively nuclear, and A3A has higher enzymatic activity that can vary from 2- to 100-fold above that of A3B depending on substrate (*e.g*., **Figure 4** and prior biochemical studies [11, 52]). It is therefore notable here that the overall genome-wide level of APOBEC3 signature mutation from A3A is only 4-fold higher than that attributable to A3B. Endogenous *A3A* and *A3B* also have both distinct and overlapping transcription programs, and both genes can be induced by a variety of conditions including viral infection and inflammation [28, 33, 35–37, 39, 42, 61, 77–82]. *In vivo*, *A3A* and *A3B* gene expression is also likely to be affected by the local tumor microenvironment, which can vary both between and within cancer types, as well as by a patient’s global state of health. Taken together with unknown and likely lengthy multi-year durations of pre-cancer and early cancer development prior to clinical manifestation, deducing the exact fractions of mutations attributable to A3A and/or A3B may be a fruitless endeavor (except in *A3B*-null tumors). Rather, it may be more prudent to focus on developing strategies to simultaneously diagnose and treat the mutagenic contributions of both enzymes.

Independent whole genome sequencing experiments have provided additional information on the APOBEC3 mutation process. Initial studies induced overexpression of A3B in 293-derived cell lines, documented the resulting DNA damage responses, and performed WGS to assess genome-wide associations [83, 84]. However, an unambiguous APOBEC3 mutation signature was difficult to extract from these whole genome sequences due to large numbers of mutations attributable to defective mismatch repair [83, 84]. A more recent study compared *de novo* mutations occurring in APOBEC3 signature positive cell lines during multiple generations of clonal outgrowth [85]. An intriguing finding from this work is that APOBEC3 signature mutations may be able to occur in an episodic manner, accumulating in some generations and not others, consistent with evidence discussed above that A3A and A3B expression can be induced by multiple signal transduction pathways. Episodic mutagenesis, however, is unexpected in cell-based systems in which continuous and relatively stochastic mutagenesis should predominate given defined media and well-controlled growth conditions. These studies were followed-up more recently by WGS comparisons of subclones of the same cancer cell lines CRISPR-engineered to lack *A3A*, *A3B*, or both genes [86]. The results of over 250 WGS combined to indicate that A3A may be the source of a significant fraction of observed APOBEC3 signature SBS mutations, A3B a smaller fraction, and another as-yet-undefined APOBEC3 enzyme an additional minor fraction. These data are complementary to the major results here, with both A3A and A3B proving capable of generating genome-wide APOBEC3 signature single base substitution mutations. Differences in the overall magnitude of A3A vs A3B mutagenesis may be due to the factors described above including differential intrinsic activity, protein expression levels, genomic DNA accessibility, cell culture conditions, and importantly durations of mutagenesis. Both studies were necessarily done in model cellular systems, each with obvious strengths, but neither capable of fully recapitulating the wide repertoire of factors that impact the actual pre- and post-transformation environments *in vivo*, including anti-tumor immune responses, which are further likely to vary between different tissue types, tumor types, and patients.

A role for A3H, haplotypes I or II, in cancer is disfavored by our results here showing that these variants are incapable of eliciting DNA damage responses or increasing the *TK* mutation frequency. Two A3H-I expressing *TK* mutant clones were subjected to WGS and no APOBEC3 mutation signature was evident. In addition, no specific evidence for A3H emerged from sequencing clonally-derived cancer cell lines [86]. However, all of these cell-based studies have limitations as discussed above and have yet to fully eliminate A3H as a source of APOBEC3 signature mutations in cancer. For instance, A3H-I may take more time to inflict detectable levels of mutation, it may be subject to different transcriptional and post-transcriptional regulatory processes, and/or it may only be mutagenic in a subset of cancer types subject to different stresses and different selective pressures.

Ultimately, the studies here showed that A3A and A3B are each individually capable of inflicting a robust APOBEC3 mutation signature in human cells and, taken together with other work summarized above, support a model in which both of these enzymes contribute to the composite APOBEC3 mutation signature reported in many different tumor types. This conclusion is supported by clinical studies implicating A3A and/or A3B in a variety of different tumor phenotypes including drug resistance/susceptibility, metastasis, and immune responsiveness [9, 20, 21, 27, 32, 42, 47, 87–93]. Thus, efforts to diagnose and treat APOBEC3 signature-positive tumors should take both enzymes into account, not simply one or the other. Such longer-term goals are not trivial given the high degree of identity between A3A and the A3B catalytic domain (>90%), the related difficulty of developing specific and versatile antibodies for detecting each enzyme, and the fact that each can be regulated differentially by a wide variety of common factors including virus infection and inflammation. Thus, we are hopeful that the HAP1-TK-M9 system, the whole genome sequences, and the A3A-specific rabbit monoclonal antibody described here will help to expedite the achievement of these goals.

## Materials and Methods

### Key resources table

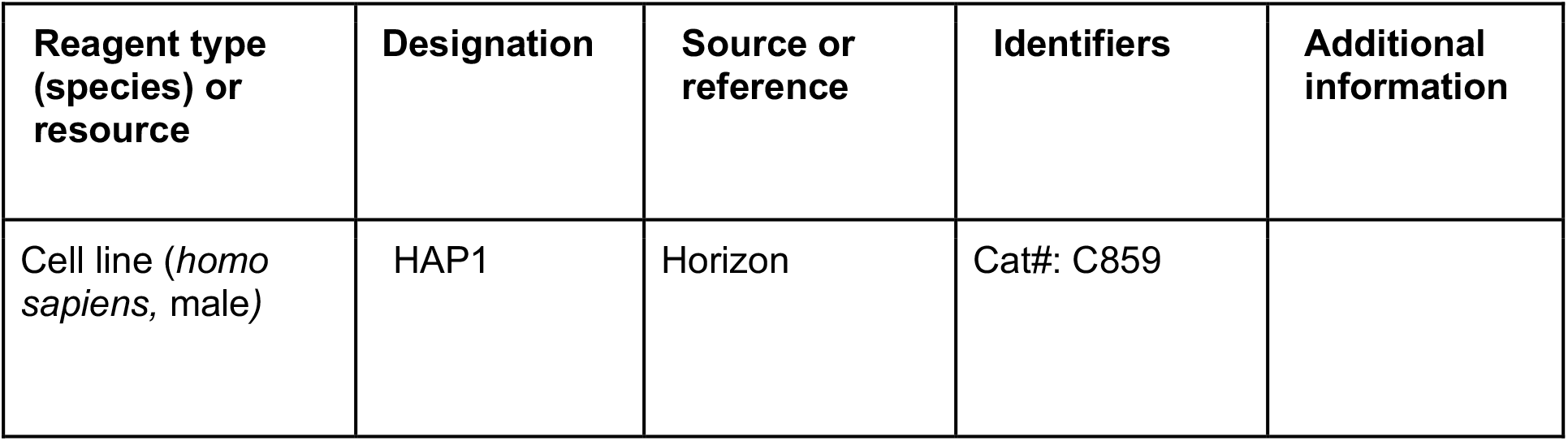
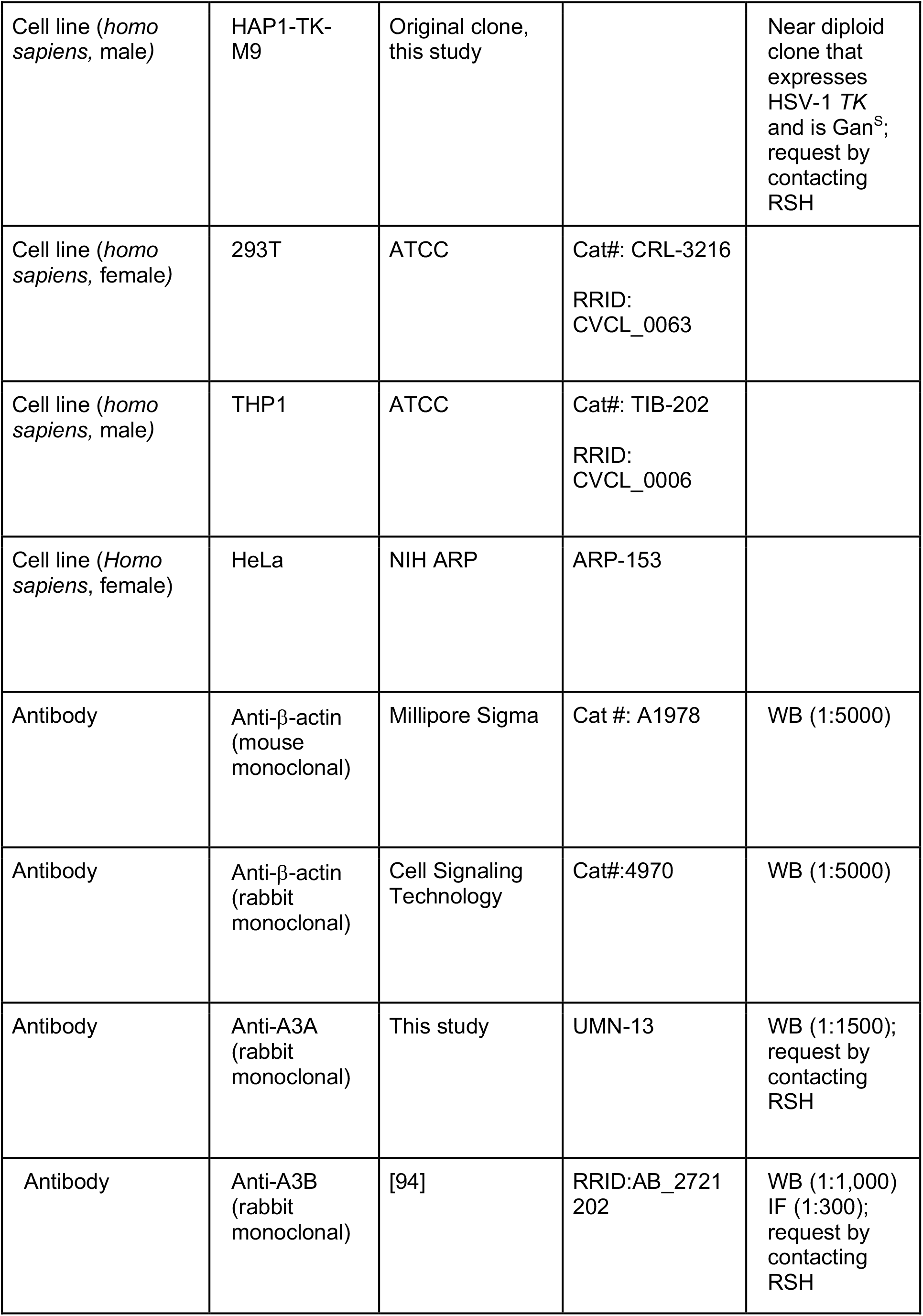
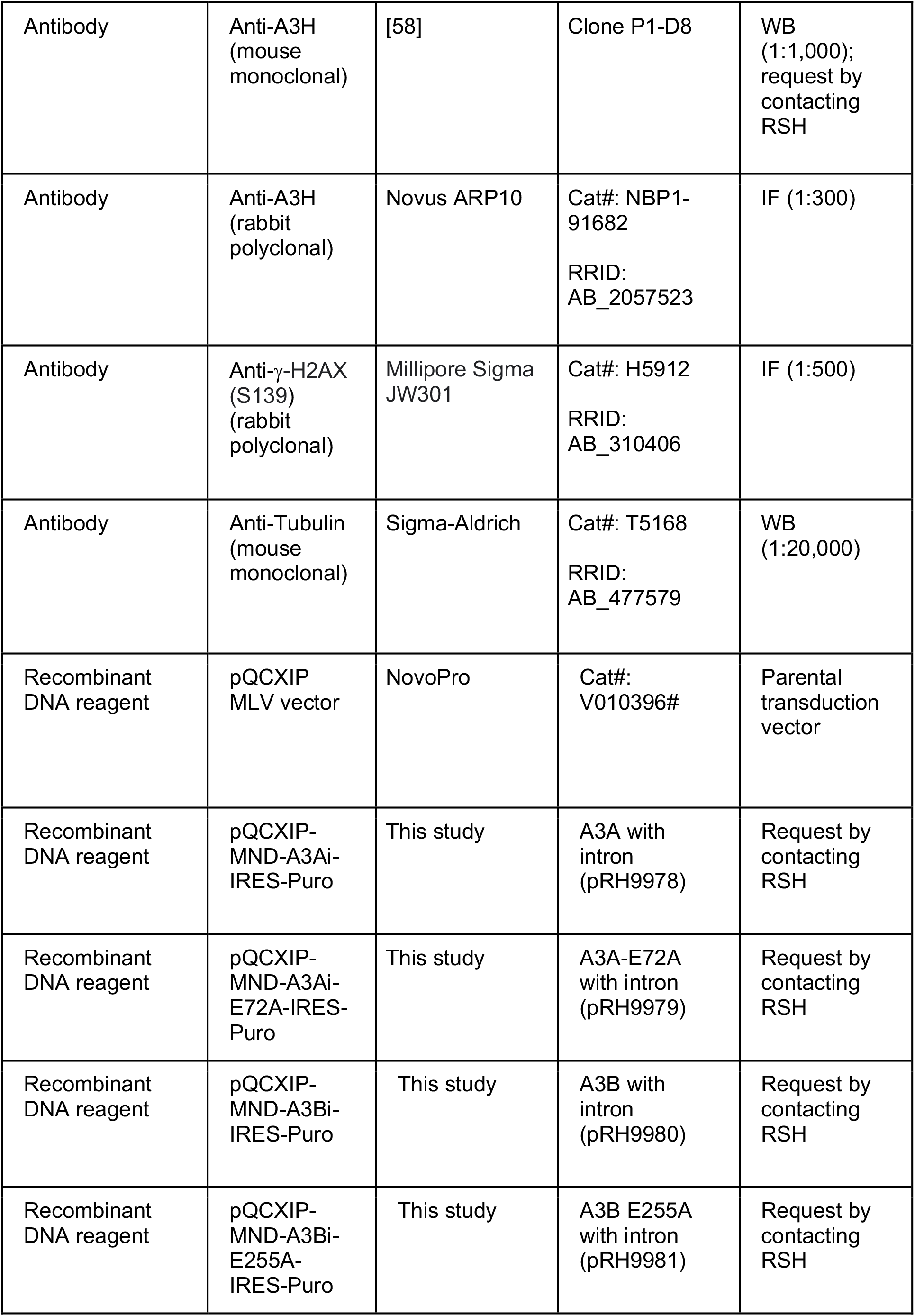
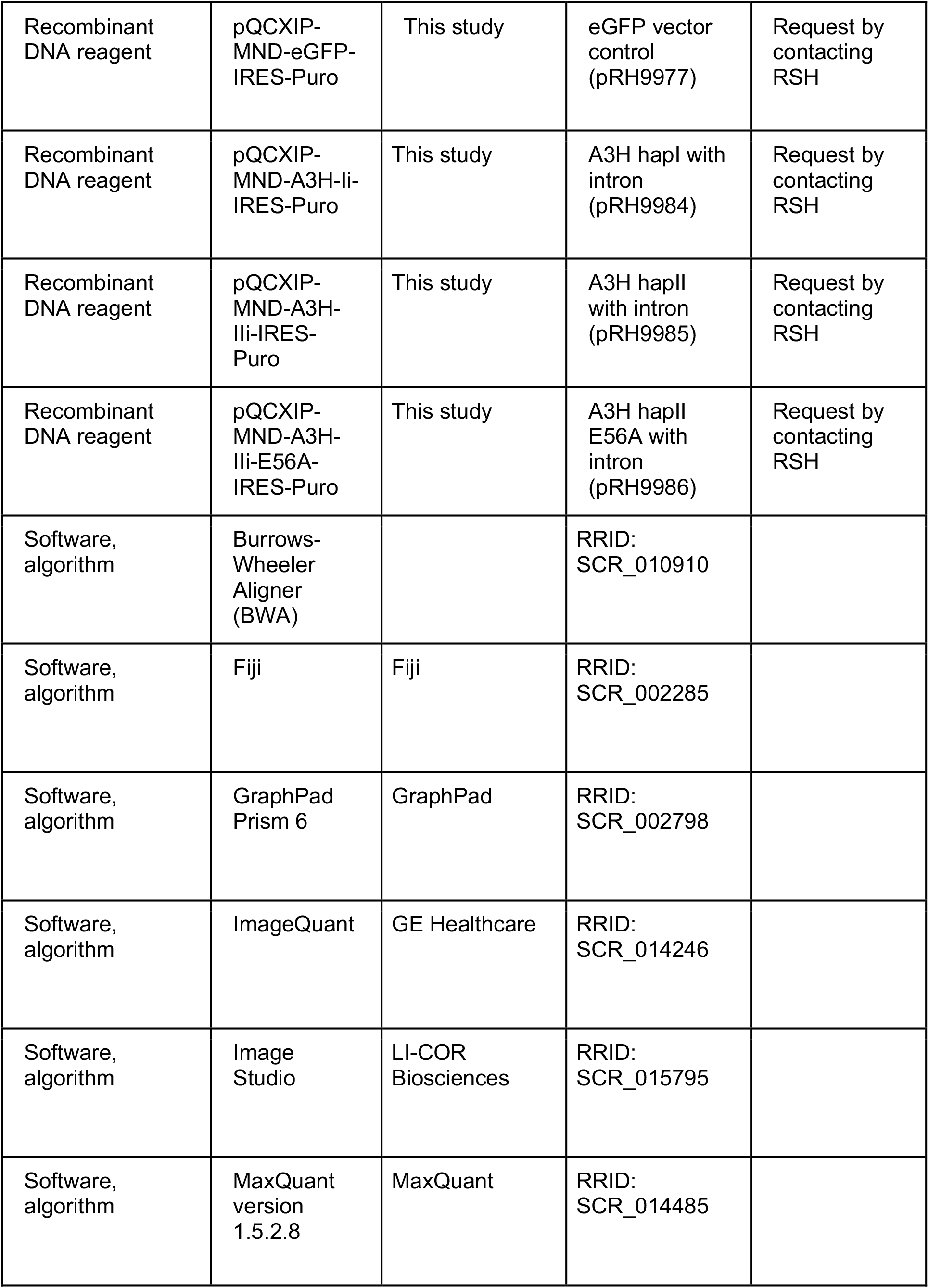
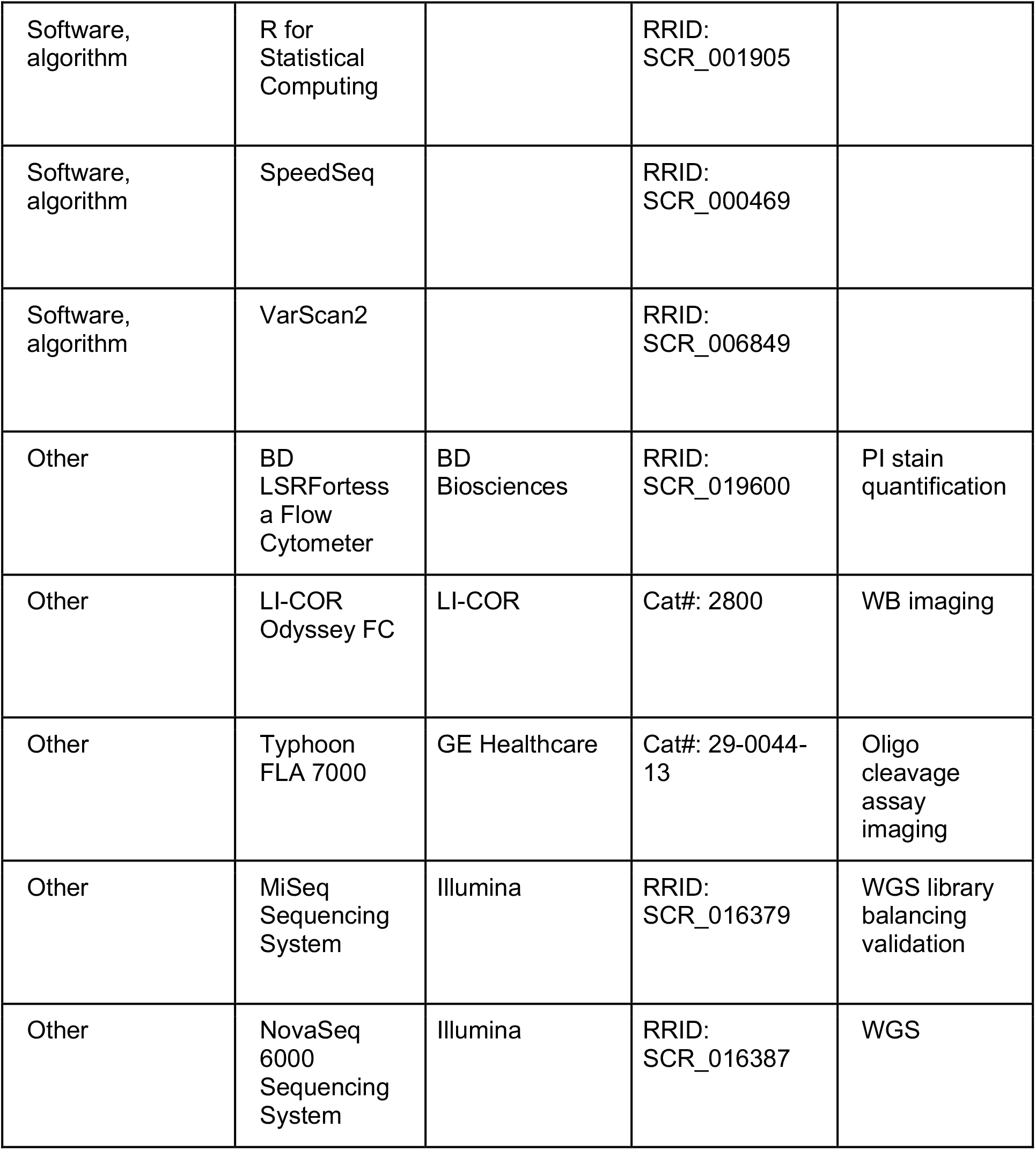

### Cell lines and culture conditions

All cell lines were cultured at 37°C under 5% CO_2_. HAP1 cells (Horizon) and derivatives (this study) were grown in IMDM (Invitrogen) supplemented with 10% fetal bovine serum (Sigma), penicillin (100 U/mL), and streptomycin (100 μg/mL). 293T and HeLa cells were cultured in DMEM (Invitrogen) with 10% fetal bovine serum (Sigma), penicillin (100 U/mL), and streptomycin (100 μg/mL). THP1 cells were cultured in RPMI (ThermoFisher) supplemented with 10% fetal bovine serum (Sigma), penicillin (100 U/mL), and streptomycin (100 μg/mL). TransIT-LT1 (Mirus) was used for all transfections. All parent and clonal lineage cell lines tested negative for mycoplasma using a PCR-based assay [95]. Puromycin (ThermoFisher) and G418 (Gold Biotechnology) were used at 1 μg/mL and 1 mg/mL, respectively. Ganciclovir (ThermoFisher) was used at 5 μM to select *TK* mutant clones.

### HAP1-TK-M9 system

The HAP1-TK-M9 system was generated by co-transfecting HAP1 parent cells with a plasmid expressing the Sleeping Beauty transposase and a separate plasmid with *TK-Neo* coding sequences flanked by SB recognition sites [9, 96]. Semi-confluent cells in 6 well plates were transfected, treated 24 hrs later with G418 (1 mg/mL), and subcloned by limiting dilution in 96 well plates to create single cell derivatives. Single cell clones were then expanded and characterized as described in the main text. The *A3H* genotype was determined by Sanger sequencing exon-specific PCR amplicons [97, 98] and further confirmed by WGS (below).

Standard molecular cloning procedures were used to create derivatives of MLV pQCXIP for expressing each A3 protein. First, pQCXIP was cut with MluI and PacI to excise the strong CMV promoter and replace it with a weaker MND promoter (a synthetic promoter containing regions of both the MLV LTR and the myeloproliferative sarcoma virus enhancer). Second, this new construct was cut with SfiI and BsiWI to insert intron-containing A3 coding sequences [67]. This was done for *A3A*, *A3B*, *A3H* haplotype-I, *A3H* haplotype-II, and appropriate catalytic mutant derivatives (E-to-A). An eGFP expressing construct was generated in parallel to use as a control in various experiments. All new constructs were confirmed by Sanger sequencing and functional assays as described in the main text.

Each MLV-based construct was co-transfected into 293T cells with appropriate packaging vectors and 48 hrs later the resulting viral supernatants were filtered (0.2 μm) and used to transduce semi-confluent HAP1-TK-M9 cells (MOI < 0.1). After 48 hrs incubation, transduced cells were selected with puromycin (1 μg/mL) and subcloned by limiting dilution to obtain A3 expressing daughter clones. Each daughter clone expressed only a single, integrated construct, which was anticipated by low MOI transduction and verified by mapping insertion sites for 5 representative daughter clones (**Figure 3 – Figure Supplement 3**). These A3 expressing and control daughter clones were expanded for 1 month and characterized as described in the main text. No overt growth/proliferation defects were noted, and all granddaughter clones expanded at similar rates. Mutation frequencies were determined by plating 5 x 10^6^ cells in 96 well flat bottom plates, treating with 5 μM ganciclovir (ThermoFisher), and after 14 days incubation counting the number of *TK* mutant colonies that survived selection. Single Gan^R^ granddaughter clones were counted using a light microscope and expanded and characterized as described in the main text including immunoblotting, DNA deaminase activity assays, *TK* sequencing, and WGS.

### Immunoblots (IB)

Cells were treated with trypsin EDTA and collected, washed in 1X PBS, and re-suspended in 100 µL of reducing sample buffer per one million cells [0.5 M Tris-HCl pH 6.8, 1% 2-mercaptoethanol, 10% sodium dodecyl sulfate (SDS), 50% glycerol]. Proteins were denatured by boiling samples for 20 min and resolved by SDS-polyacrylamide gel electrophoresis. Proteins were transferred to a PVDF-FL membrane (Millipore Sigma) and blocked in 5% milk in 1X PBS. Primary and secondary antibodies were incubated in blocking buffer, with the addition of 0.2% SDS for fluorescent antibodies. The primary antibodies used were anti-A3A (UMN-13, IB: 1:1500, IF: 1:300 [this study]), anti-A3A/B (5210-87-13, IB: 1:1,000, IF: 1:300 [94]), anti-A3H (P1-D8, IB: 1:300 [58]), anti-Tubulin (Sigma Aldrich, 1:20,000), and anti-ý-Actin (Millipore Sigma, mouse mAb, 1:5000 or Cell Signaling Technology, rabbit mAb, 1:5000, as indicated). The secondary antibodies used were anti-rabbit HRP-linked (CST 7074, 1:2,000), IRDye 800CW goat anti-mouse (LI-COR Biosciences, 1:10,000), Alexa Fluor 680 goat anti-rabbit (Molecular Probes, Eugene, OR, USA, 1:10,000), and Alexa Fluor 680 goat anti-mouse (Molecular Probes, Eugene, OR, USA, 1:10,000). Membranes were imaged using a LI-COR Odyssey instrument or LI-COR Odyssey-Fc instrument for HRP visualization (LI-COR Biosciences).

### DNA deaminase activity assays

Whole cell extract (WCE) assays: ssDNA deamination activities were measured using WCE prepared using 100 μL HED lysis buffer per 1 million cells (25 mM HEPES, 15 mM EDTA, 10% Glycerol, 1 mM DTT, and 1 protease inhibitor tablet [Roche]). Samples were sonicated in a water bath sonicator to ensure complete lysis. A3-containing lysates were incubated at 37°C for 1 hr (A3A and A3B) or 4 hrs (A3H) with purified human UNG2 and a ssDNA substrate containing either a single TCA or a single TCT trinucleotide motif (5’-ATTATTATTATTCAAATGGATTTATTTATTTATTTATTTATTT-FAM [A3A and A3B]; 5’-ATTATTATTATTCTAATGGATTTATTTATTTATTTATTTATTT-FAM [A3H]) following established protocols [9, 99]. After this initial incubation, the reaction was treated with 100 mM NaOH for 5 min at 95°C. The reaction was run out on a 15% TBE-urea acrylamide gel to separate substrate oligo from cleaved product oligo and imaged on a Typhoon FLA 7000 with ImageQuant TL 8.2.0 software (GE Healthcare).

Recombinant enzyme assays: A3A- and A3B-mycHis were prepared from transfected 293T cells as reported [14, 16, 100–102]. Single hit kinetics were ensured by incubating 25 nM of each protein with 800 nM substrate in reaction buffer (25 mM HEPES pH 7.4, 50 mM NaCl, 5 mM imidazole) for the indicated times at 37°C. Reactions were stopped by freezing in liquid nitrogen and then were heated to 95°C to denature the enzymes. Reactions were then treated with 0.5 U/reaction uracil DNA glycosylase (NEB, USA) for 10 min at 37°C. The resulting abasic sites were cleaved by incubation with 100 mM NaOH and heating to 95°C for 5 min. Products were separated by 20% TBE-Urea PAGE, imaged on a Typhoon FLA-7000 (GE Healthcare, USA), and quantified using ImageQuant TL 8.2.0 software (GE Healthcare). Deamination of the target cytosine was calculated by dividing the total reaction product by the total amount of starting substrate. The oligonucleotide substrates were analyzed by both denaturing and native 20% PAGE to determine the extent of hairpin formation. The oligos (and a previously reported NUP93-noHP oligo) were heated to 70°C and slowly cooled to 37°C in HEPES buffer as above. 1 pmol of each oligo was then mixed with agarose gel loading dye (30% Ficoll 400 in 1x TAE, xylene cyanol, bromophenol blue) or with DNA PAGE loading dye (80% formamide in 1x TBE, xylene cyanol, bromophenol blue) and separated by native and denaturing PAGE, respectively.

### DNA content by flow cytometry

Propidium iodide (PI) staining was used to assess the ploidy of HAP1 clones relative to THP1 (ATCC, Cat#: TIB-202) as a confirmed diploid control [103]. Cells were trypsinized and suspended in 100 μL of 1X PBS per 1 million cells. Then, 500 μL ice-cold ethanol was added to cell suspensions and incubated at -20°C for 1 hr to fix the cells. After fixation, cells were pelleted and washed in 1X PBS. Cells were finally suspended in 500 μL of FxCycle PI stain (Invitrogen) and incubated for 30 min at RT in the dark to stain the cells. Cells were spun down and resuspended in 300 μL of the PI stain solution and placed in a 96 well round bottom plate for flow cytometry analysis using a BD LSRFortessa flow cytometer (with high-throughput 96 well adapter system). A minimum of ten thousand events were acquired for each condition.

### RT-qPCR

Total RNA was extracted using the High Pure RNA isolation kit (Roche). cDNA was synthesized using SuperScript First-Strand RT (ThermoFisher). Quantification of mRNA was done using validated primer sets for all human *A3* genes relative to the housekeeping gene *TBP* [9, 78, 96, 97]. All RT-qPCR reaction were performed using SsoFast SYBR Green mastermix (Bio-Rad) in 384-well plates on a LightCycler 480 (Roche) following the manufacturer’s protocol. Statistical analyses were done using GraphPad Prism 6 and R.

### Immunofluorescent (IF) microscopy

IF microscopy was conducted as described [104, 105]. As a positive control for DNA damage, 4 μM cisplatin (*cis*-diamminedichloroplatinum II, Selleck Chemical) was incubated with cells for 24 hrs. Cells were grown at low density in 4-chamber, tissue culture-treated glass slides (Falcon) prior to fixation. The cells were then fixed with 4% paraformaldehyde in PBS for 15 min at room temperature. Permeabilization followed using PBS containing 0.2% Triton X-100 (Sigma Aldrich), before rinsing with PBS. The cells were blocked using an IF blocking solution (0.1% triton X-100, 5% goat serum in PBS) for 1 hr at room temperature, then incubated with primary antibody overnight at 4°C. The primary antibodies used were anti-A3A/B/G (5210-87-13, 1:300) [94], anti-A3A (UMN-13, 1:300 [this study]), anti-A3H (Novus, 1:300) [58], and anti-γ-H2AX (JBW301, Millipore Sigma, 1:500). Following this, cells were washed with PBS and incubated with a fluorophore-conjugated secondary antibody for 1 hr at room temperature in the dark. The secondary antibodies used were Alexa Fluor 488 goat anti-mouse (Invitrogen 1:1,000) and Alexa Fluor 594 goat anti-rabbit (Invitrogen, 1:1,000). Both primary and secondary antibodies were diluted in blocking buffer. Hoechst 33342 (Mirus) was used at a final concentration of 1 μg/mL to stain nuclei for another 15 min, before the slides were washed three times with PBS and mounted using antifade mounting media (Cell Signaling 9071). Images were captured at 60x magnification using a Nikon ECLIPSE Ti2 microscope. The number of γ-H2AX foci per cell were counted for at least 50 cells per condition for each experiment. The nuclear intensity of the γ-H2AX signal and the cellular intensity of each A3 were measured by ImageJ2 software (2.9.0/1.53t). Statistical analyses were conducted in GraphPad Prism 9.

### Alkaline comet assays

HeLa cells transfected with individual A3 or control constructs were harvested 24 hrs post-transfection and resuspended in ice-cold 1X phosphate-buffered saline (PBS, Ca^2+^ and Mg^2+^ free, at a density of 10^5^ cells/mL). As a positive control, mock-transfected cells were treated with 2 μM camptothecin (CPT, Millipore Sigma) for 2 hrs prior to harvesting as above. The CometAssay ESII kit was used for all alkaline Comet assays, following the manufacturer’s protocol (BioTechne). Cells were resuspended in low-melt agarose and spread at low density on a glass slide to secure the cells in place for lysis. DNA unwinding and electrophoresis were performed according to the manufacturer’s alkaline protocol. Comet tail moments were measured for at least 50 cells per condition using the OpenComet plugin for Image J [106]. Statistical analyses were done using GraphPad Prism 9.

### *TK* sequencing

Cells were harvested and genomic DNA was isolated using the Puregene DNA isolation protocol. The *TK* cassette was amplified from genomic DNA using 5’-ATCTTGGTGGCGTGAAACTC and 5’-CTTCCGGTATTGTCTCCTTCC. PCR products were cleaned-up using the GeneJet Gel Extraction Kit (Thermo Scientific) and Sanger sequenced with 4 different primers to cover the full open reading frame (5’-ATCTTGGTGGCGTGAAACTC, 5’-GGTCATGCTGCCCATAAGGTA, 5’-CCGTTCTGGCTCCTCATATC, and 5’-CTTCCGGTATTGTCTCCTTCC.

### Whole genome sequencing (WGS) and analyses

Genomic DNA was prepared from cell pellets (1 million cells) using Allprep DNA/RNA mini kit (Qiagen). Whole genome libraries were sequenced 150x2 bp on a NovaSeq 6000 (Illumina) to a target read depth of 30X coverage for all granddaughter clones as well as the parental HAP1-TK-M9 mother clone. Resulting sequences were aligned to the human genome (hg38) using SpeedSeq [107], which relies on the Burrows-Wheeler Aligner, BWA (version 0.7.17). PCR duplicates were removed using Picard (version 2.18.16). Reads were locally realigned around INDELs using GATK3 (version 3.6.0) tools RealignerTargetCreator to create intervals, followed by IndelRealigner on the aligned bam files. Single base substitutions and small INDELs were called in each clone relative to the bam file generated from the HAP1-TK-M9 mother clone using Mutect2 from GATK3 (version 3.6.0). SBSs that passed the internal GATK3 filter with minimum 4 reads supporting each variant, minimum 20 total reads at each variant site and a variant allele frequency over 0.05 were used for downstream analysis. SBSs were analyzed in R (version 4.0.5) using the MutationalPatterns [108] and deconstructSigs R packages (version 1.8.0 [109]). All visualizations were generated using the ggplot2 package (version 3.3.5). The indel landscapes were generated using the MutationalPatterns R package [108] following PCAWG definitions [1]. All individual clone data from each condition were pooled for presentation.

COSMIC single base substitution mutation signatures (v3 – May 2019 https://cancer.sanger.ac.uk/cosmic/signatures/SBS/) were obtained from https://www.synapse.org/#!Synapse:syn11738319. *De-novo* non-negative matrix factorization of mutational signatures was performed with the “extract_signature” command from the MutationalPatterns package, with a rank of 2 and 100 iterations. TCW mutation enrichment scores were calculated as described [50, 65]. Sequence logos of -2 to +2 sequence surrounding C-to-T mutations were created using the ggseqlogo (version 0.1) package. Putative hairpin structures were predicted using the nBMST tool [110] and human genome GRCH38 with a minimum stem length of 6 bases. The loop criteria established by Buisson *et al*. with ssDNA loop lengths of 3 to 11 nucleotides were used to search for APOBEC3 signature mutations in these regions [52].

### APOBEC expression and mutation signature analyses in TCGA and ICGC data sets

TCGA primary breast tumors represented by both RNA-seq and whole exome sequencing were downloaded from the Firehose GDAC resource through the Broad Institute pipeline (http://gdac.broadinstitute.org/) for multiple tumor tissue types. APOBEC3 mutation signatures were determined as described [5, 65] using the deconstructSigs R package [109]. APOBEC3 mutation enrichment scores were calculated using the hg19 reference genome and published methods [50]. Enrichment score significance was assessed using a Fisher exact test with Benjamini-Hochberg false discovery rate (FDR) correction. All data analyses and visualizations were conducted using R and the ggplot2 package (https://www.R-project.org/).

### *TK* integration site determination

To determine the integration site of the single copy *TK-Neo* construct, a *TK* reference sequence was provided as an additional chromosome during alignment of the WGS reads to the reference genome (hg38). Reads that mapped to this region were then categorized as discordant and realigned to hg38 using GRIDSS (v2.2) [111] to determine the site of integration.

### Clustering analysis

All primary breast tumor whole genome sequencing variant information from International Cancer Genome Consortium (ICGC) was downloaded from the ICGC data portal (https://dcc.icgc.org). SBSs used in these analyses included only C-to-T variants in TC dinucleotide contexts (TCA, TCC, and TCT) and excluded all mutations in CG motifs due to potential overlap with spontaneous water-mediated methyl-C deamination. SBSs meeting these inclusion criteria from all clones expressing A3A, A3B, A3A-E72A, A3B-E255A, and A3H-I were pooled per condition for this analysis. A matrix comprised of the number of mutations within a pentanucleotide across all samples within a cancer type was generated, and counts were normalized to frequency within each cancer type. The resulting matrix was then clustered using the hclust function in R with the classical Euclidean distance as the distance method for clustering, which was then plotted as dendrograms. Mutation signatures were calculated using deconstructSigs as described above.

## Data and code availability

All HAP1-TK-M9 clone alignment files (FASTQ and BAM format) are available through the Sequence Read Archive under the BioProject accession number PRJNA832427.

## Author contributions

R.S. Harris conceptualized and designed the overall project. M.A. Carpenter, M.A. Ibrahim, M.C. Jarvis, M.R. Brown, P.P. Argyris, and W.L. Brown performed experiments. M.C. Jarvis, M.A. Carpenter, M.A. Ibrahim, N.A. Temiz, M.R. Brown, P.P. Argyris, and W.L. Brown did formal data analysis. R.S. Harris, M.C. Jarvis, and D. Yee contributed to funding acquisition. R.S. Harris and M.C. Jarvis drafted the manuscript, and all authors except M.C. Jarvis contributed to manuscript proofing and revision.

## Additional contributions

We thank Justin Leung, Eloïse Dray, Weixing Zhao, and Wenjing Li for expert assistance with IF microscopy and comet assays, Arad Moghadasi and Sofia Moraes for initial technical support, and Kyle Richards for helping to set-up spectral karyotype analysis. We also thank the UMN Cytogenetics Core for spectral karyotyping and Scott McIvor for sharing a plasmid construct with the *MND* promoter. The results presented here are in part based upon data generated by the TCGA Research Network: http://www.cancer.gov/tcga. Additionally, data from the International Cancer Genome Consortium (ICGC) were used in these analyses: https://dcc.icgc.org/.

## Figure Legends – Main and Figure Supplements

**Figure 1 – Figure Supplement 1.**
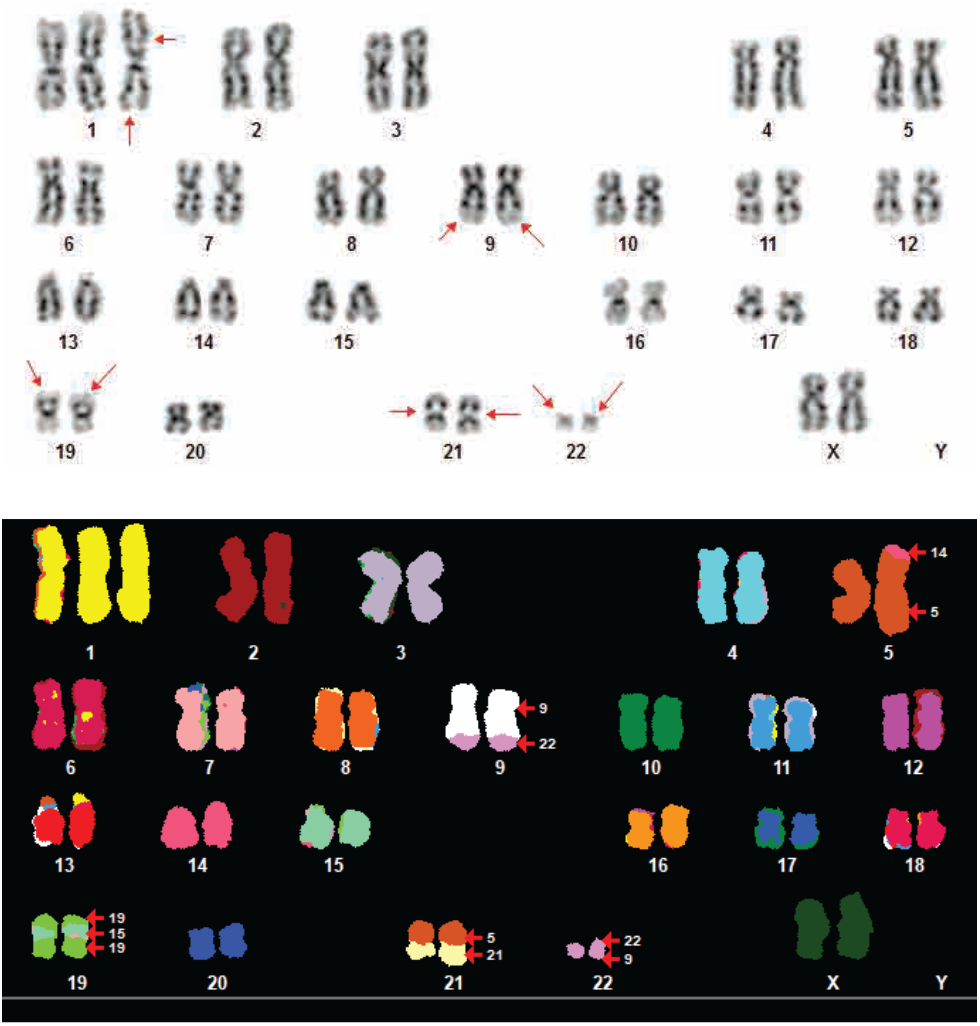
HAP1-TK-M9 karyotype analysis. Representative G-band and spectral karyotype (SKY) images of HAP1-TK-M9 M-phase chromosomes showing a near diploid DNA content and previously reported aberrations including the reciprocal chromosome 9:22 translocation (Philadelphia chromosome) characteristic of CML tumor cells. The Y-chromosome is missing, as reported for the KBM7 parent line of HAP1.

**Figure 1 – Figure Supplement 2.**
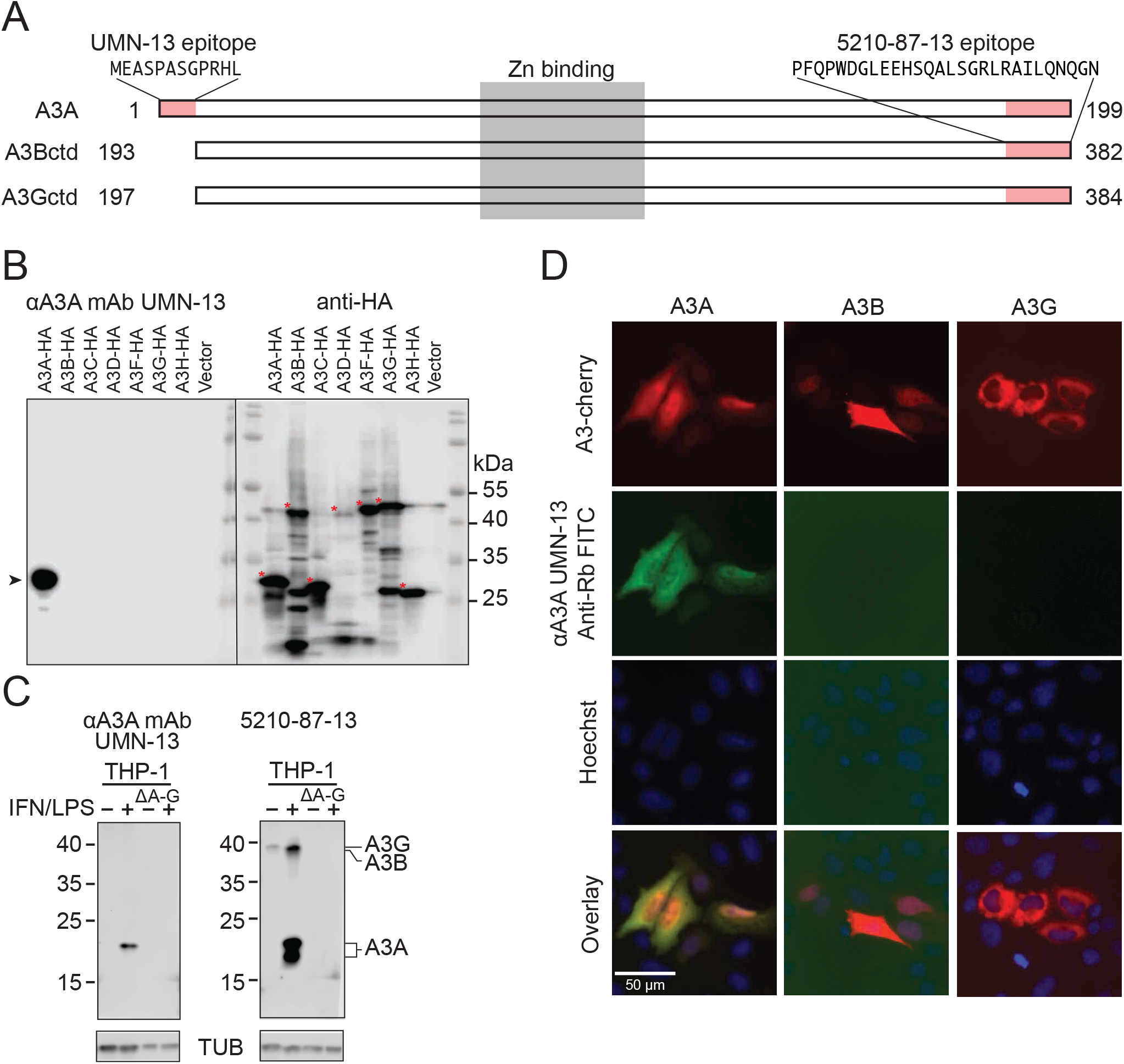
Validation of UMN-13 as a custom rabbit anti-human APOBEC3A mAb. (**A**) Schematic of human A3A, A3B, and A3G indicating the unique N-terminal epitope used here to generate the A3A-specific mAb UMN-13. The schematic also shows the C-terminal epitope used previously to generate the versatile 5210-87-13 mAb that recognizes these three enzymes. (**B**) Comparative immunoblots of whole cell extracts from 293T cells expressing each of the 7 human A3 family members with C-terminal HA tags. The blot was probed first with our custom rabbit anti-human A3A mAb UMN-13 (left) and, after stripping, a commercial anti-HA mAb as an expression control (right). The positions of the full-length proteins are indicated by red asterisks. (**C**) Comparative immunoblots of whole cell extracts from the monocytic cell line THP-1 and a clonal derivative lacking *A3A*-through-*A3G* (*ΔA-G*), each treated with DMSO as a control or LPS/IFN-α to induce expression of multiple A3s including *A3A* and *A3G*. The UMN-13 mAb blot on the left shows a single band representing full-length A3A (starting at Met1), which is absent in the deletion mutant, and the 5210-87-13 mAb blot on the right shows A3G (strong top band), A3B (weak band just below A3G), and both A3A translation products (strong band for full-length A3A starting at Met1 and a faster-migrating band for the shorter isoform starting at Met13), which are all absent in the deletion mutant. (**D**) IF microscopy images of 293T cells expressing A3A-mCherry, A3B-mCherry, or A3G-mCherry. Only the A3A construct is detected by the UMN-13 mAb as indicated by green signal in the same cells and cellular compartments as the A3A-mCherry signal.

**Figure 1 – Figure Supplement 3.**
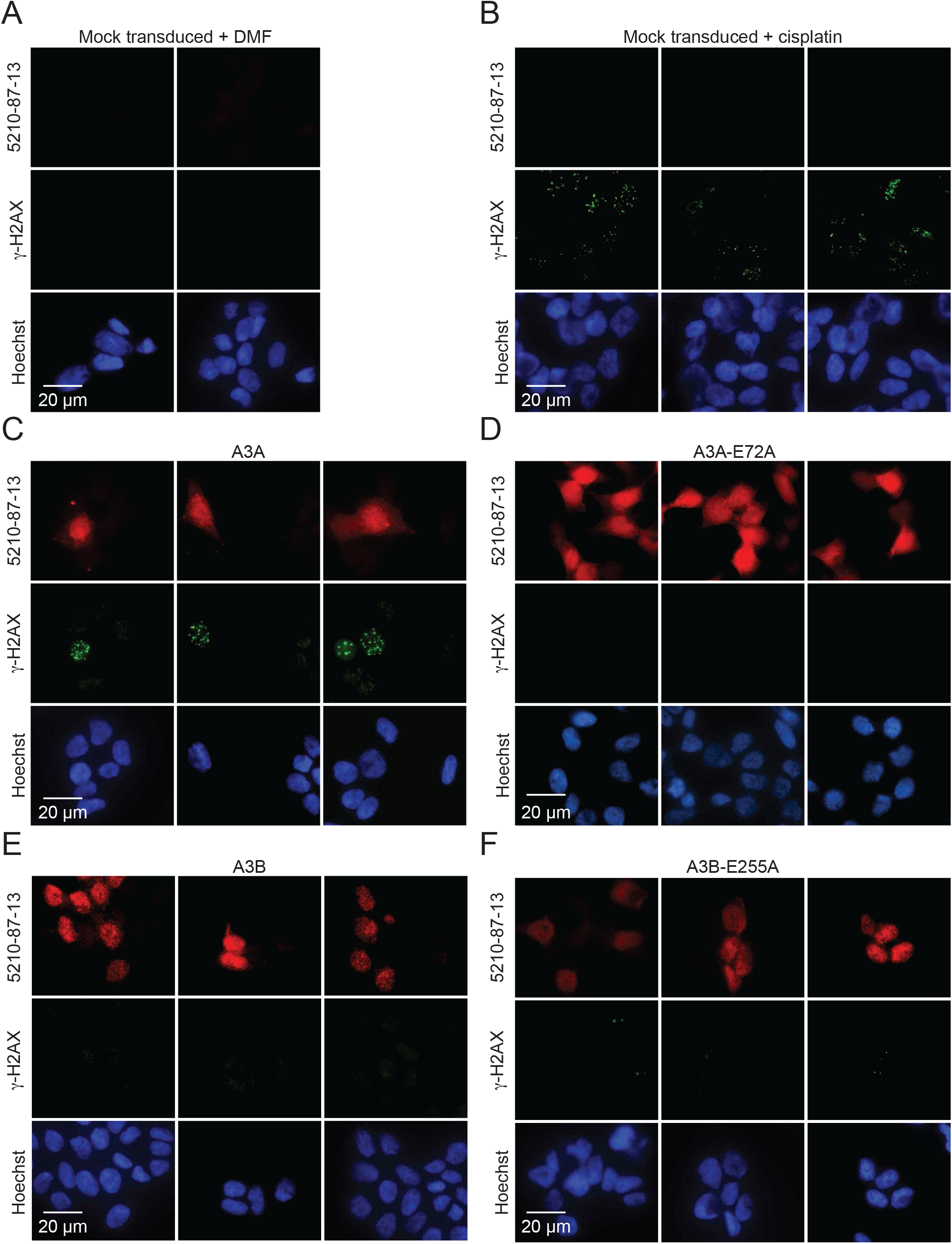
Additional IF microscopy images of HAP1-TK-M9 cells expressing A3A, A3B, and catalytic mutant derivatives. (**A-B**) Additional IF-images of HAP1-TK-M9 cells mock-transduced and DMF or cisplatin treated, respectively, and stained as indicated in parallel with cells in panels C-F (scale = 20 µm). (**C-F**) Additional IF-images of HAP1-TK-M9 cells transduced with the indicated A3 expression constructs and stained with the 5210-87-13 rabbit anti-human A3A/B mAb, γ-H2AX, and Hoechst (scale = 20 µm).

**Figure 1 – Figure Supplement 4.**
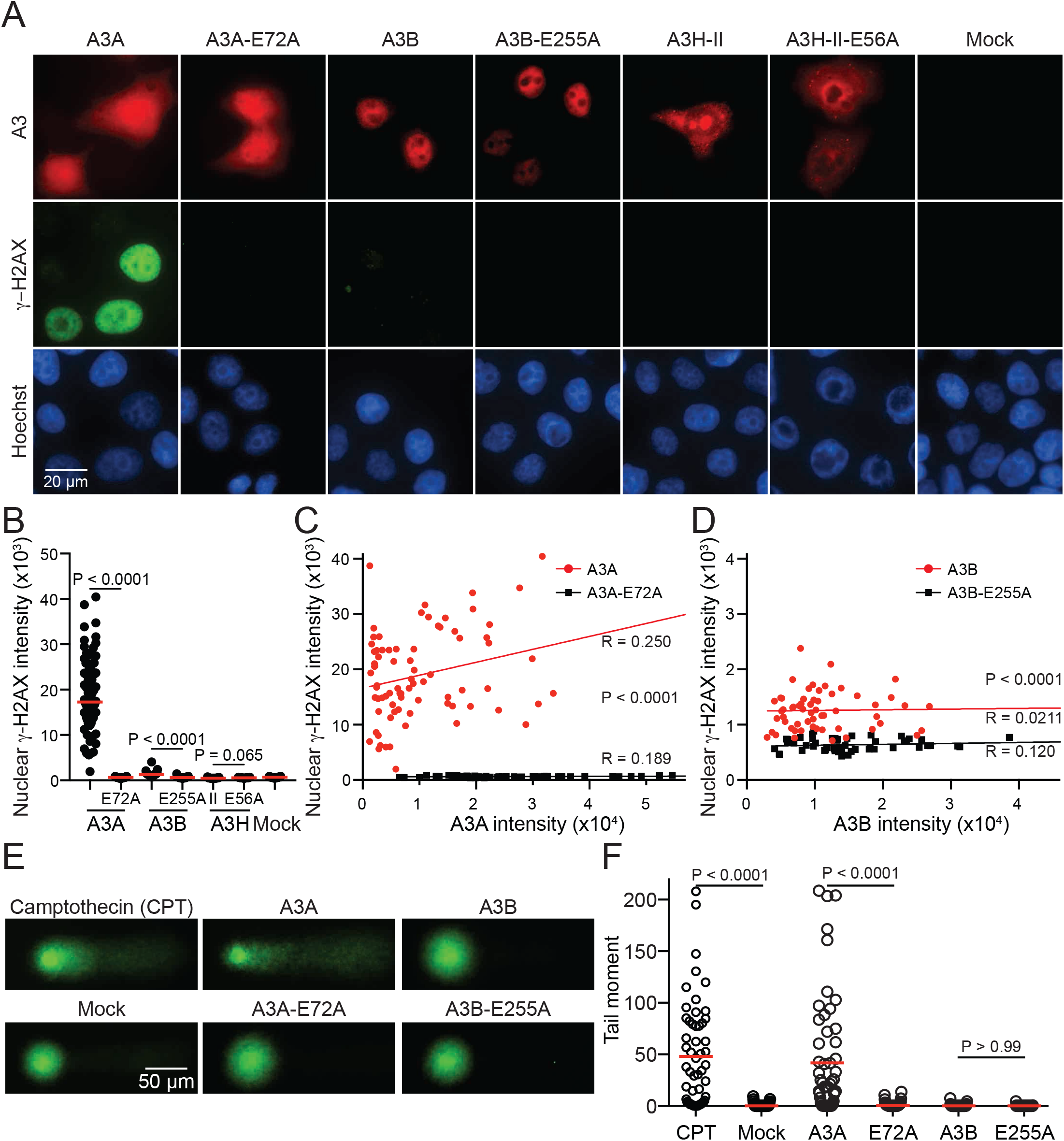
A3 localization, γ-H2AX staining, and DNA breakage in HeLa cells. (**A**) IF-images of HeLa cells transfected with the indicated A3 expression constructs and stained for A3A/B (5210-87-13 rabbit anti-human A3A/B mAb) or A3H (Novus ARP10 rabbit anti-human A3H pAb), γ-H2AX, and Hoechst (scale = 20 µm). (**B**) Quantification of nuclear γ-H2AX staining intensity in the different A3 expressing conditions in panel A (n>50 cells per condition; red bars indicate mean expression levels; p-values by Welch’s t-test). (**C**) Dot plot of nuclear γ-H2AX staining intensity versus A3A staining levels for A3A and A3A-E72A expressing cells shown in panel B. Correlation coefficients (R-values) indicate that A3A expression levels fail to correlate with nuclear γ-H2AX staining intensity. The p-value by Welch’s t-test reflects the difference between that A3A and A3A-E72A data sets. (**D**) Dot plot of nuclear γ-H2AX staining intensity versus A3B staining levels for A3B and A3B-E255A expressing cells shown in panel B. Correlation coefficients (R-values) indicate that A3B but not A3B-E255A expression levels associate with nuclear γ-H2AX staining intensity. The p-value by Welch’s t-test reflects the difference between that A3B and A3B-E255A data sets. (**E**) Representative comets from HeLa cells mock transduced, treated with 2 μM camptothecin (CPT), or transduced with expression constructs for A3A, A3B, or catalytic mutant derivatives (scale = 50 μm). (F) Quantification of tail moment for >50 cells per condition indicated in panel E (red bars indicate mean tail moment; p-values by one-way ANOVA).

**Figure 2 – Figure Supplement 1.**
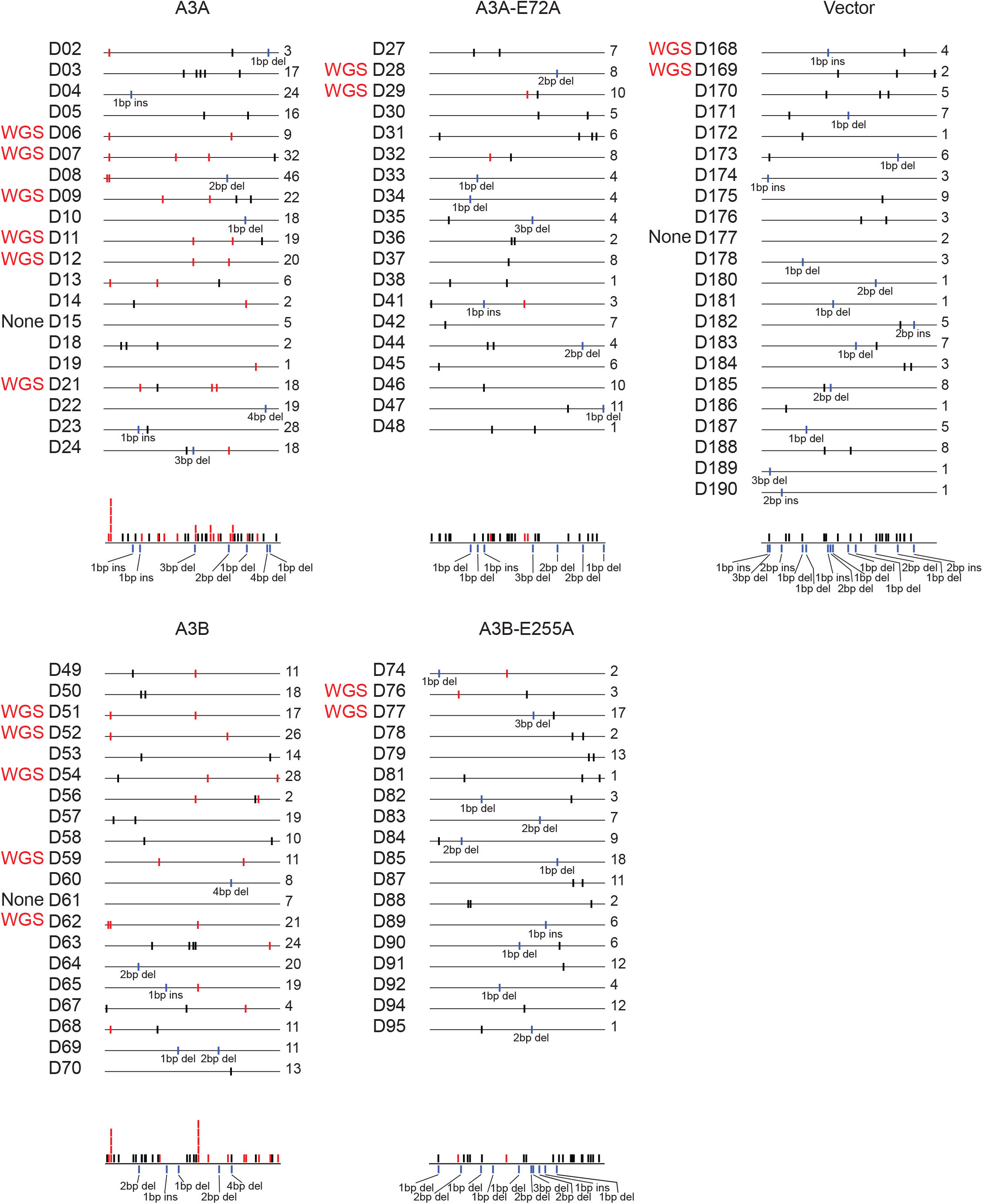
Mutations in *TK* genes derived from ganciclovir-resistant clones. Schematics of individual *TK* mutations in ganciclovir-resistant clones. T[C>G/T]W mutations are shown in red, other SBSs in black, and INDELs in blue. Composite mutation schematics are shown below for each condition.

**Figure 3 – Figure Supplement 1.**
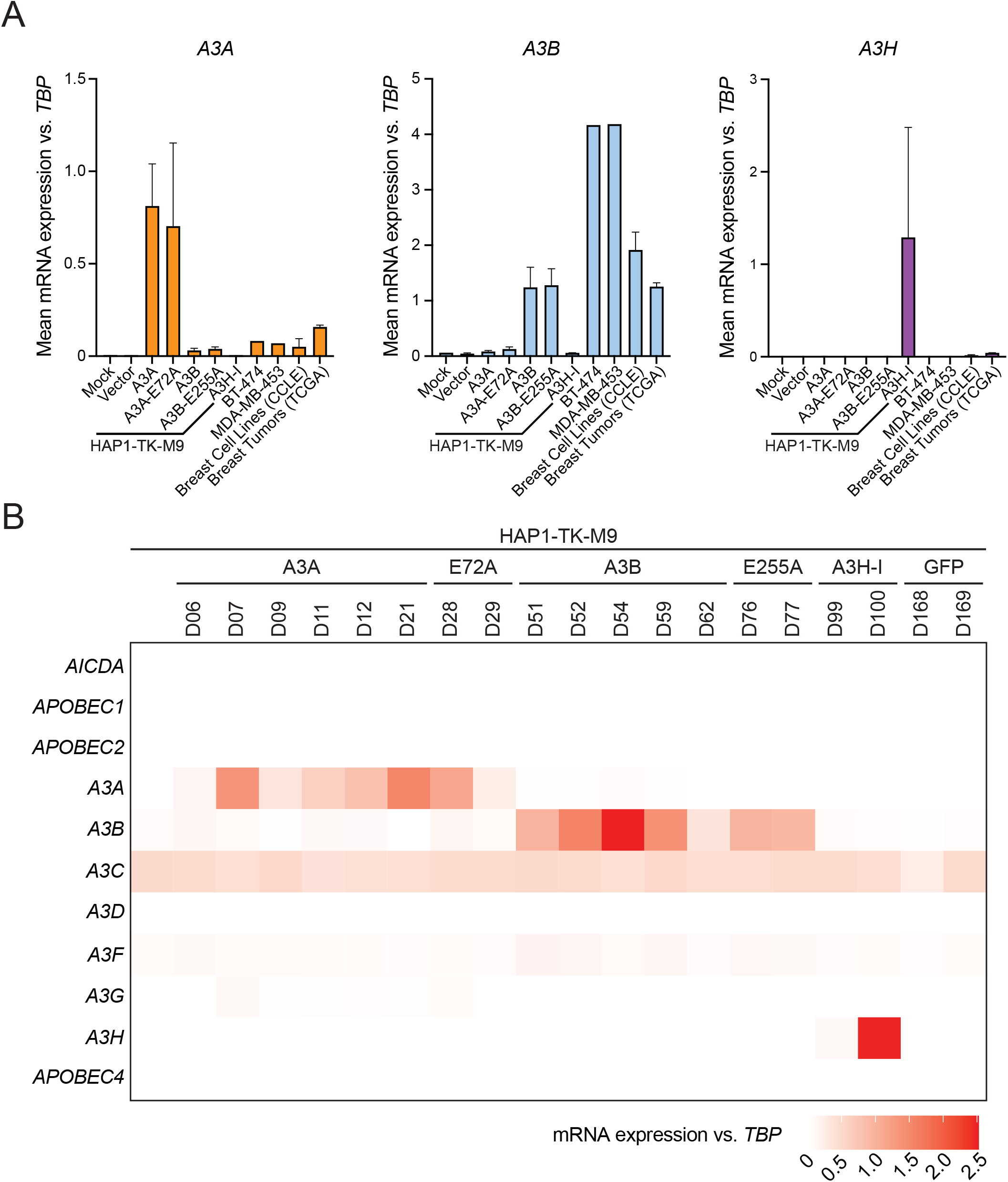
*A3* mRNA expression in the HAP1-TK-M9 system. (**A**) *A3A*, *A3B*, and *A3H* mRNA expression levels relative to those of the housekeeping gene *TBP* for the indicated HAP1-TK-M9 conditions (RNA-seq FKPM from n≥2 Gan^R^ clones for each condition; mean +/- SD shown). RNA-seq data from A3 signature-high breast cancer cell lines (BT-474 and MDA-MB-453), CCLE breast-derived cell lines (n=52), and TGCA primary breast cancers (n=1093) for comparison (mean +/- SD). (**B**) A heatmap depicting mean expression levels of all 7 human *APOBEC3* family members, in addition to *AICDA*, *APOBEC1*, *APOBEC2*, and *APOBEC4*, relative to those of the housekeeping gene *TBP* (RNA-seq values are FKPM; n≥2 for each condition to provide matching data sets for the Gan^R^ clones subjected to WGS). Endogenous *A3C* provides a consistent internal control.

**Figure 3 – Figure Supplement 2.**
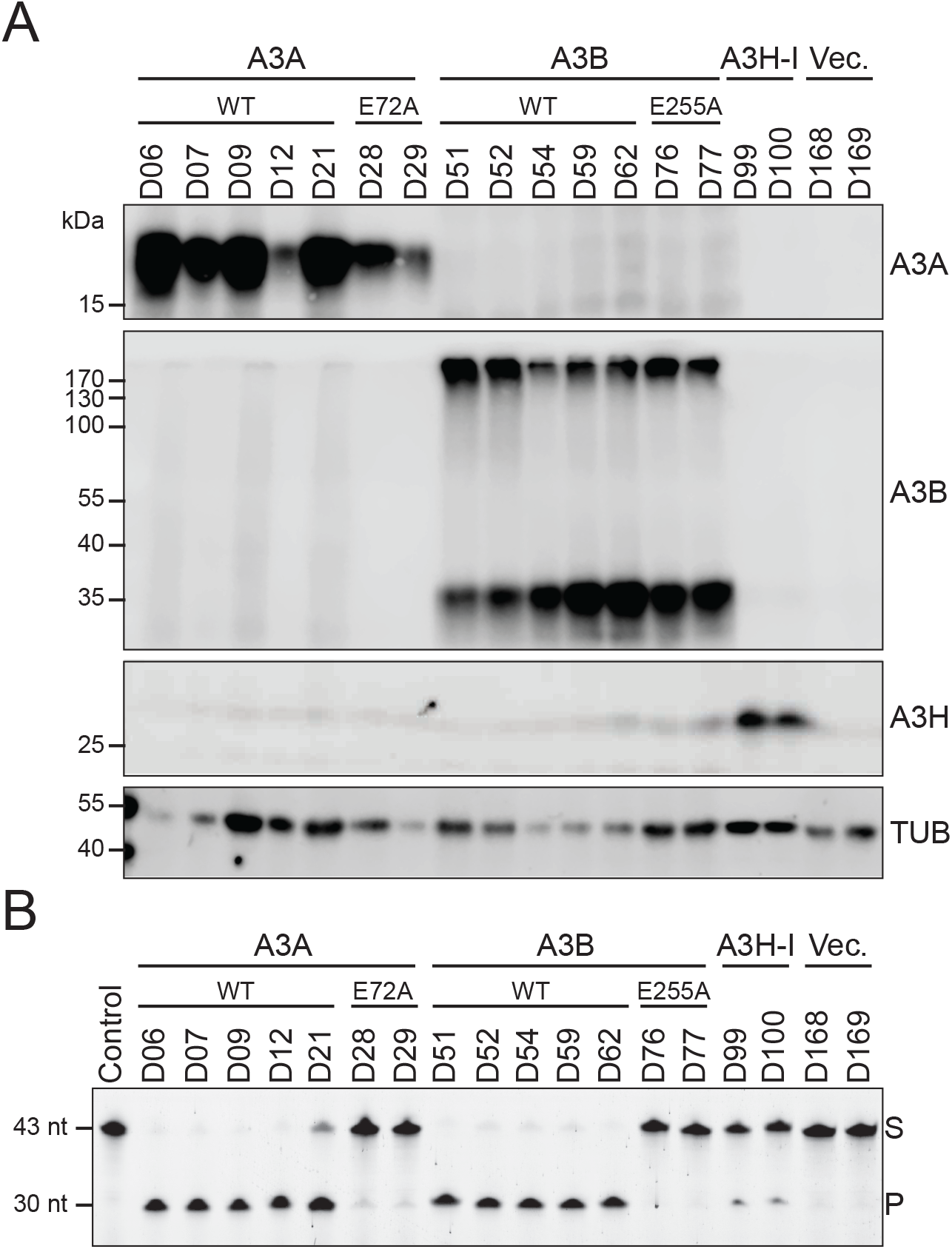
A3 protein expression in Gan^R^ clones. (**A**) Immunoblots of A3A, A3B, and A3H in the indicated Gan^R^ clones. Tubulin (TUB) is a loading control. (**B**) Deaminase activity of WCE on ssDNA from the same clones (S, substrate; P, product).

**Figure 3 – Figure Supplement 3.**
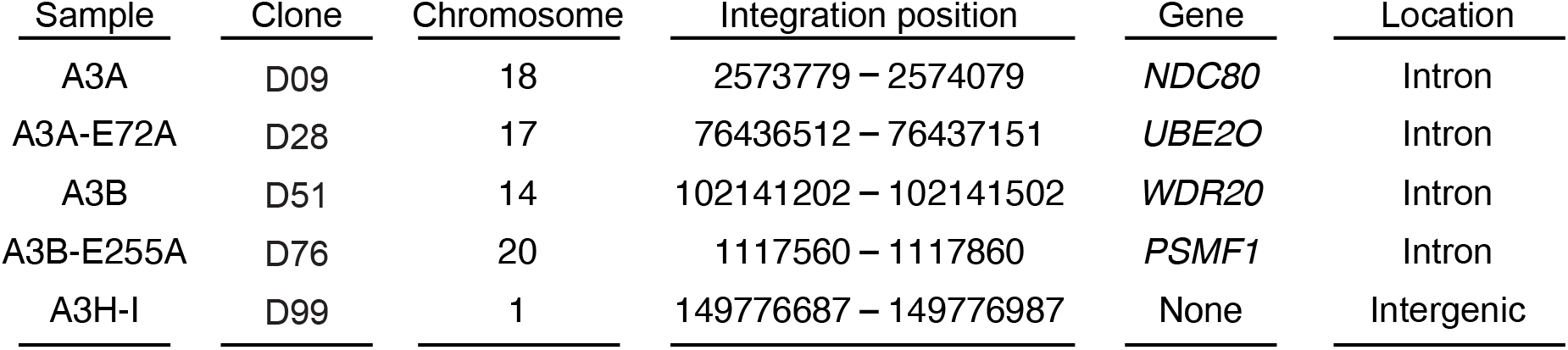
Representative MLV-A3 insertion sites in HAP1-TK-M9 clones. A table indicating chromosomal locations of representative MLV-A3 insertion sites in HAP-1-TK-M9 clones. In each clone, a single MLV-A3 insertion is positioned in the window between the indicated nucleotides.

**Figure 3 – Figure Supplement 4.**
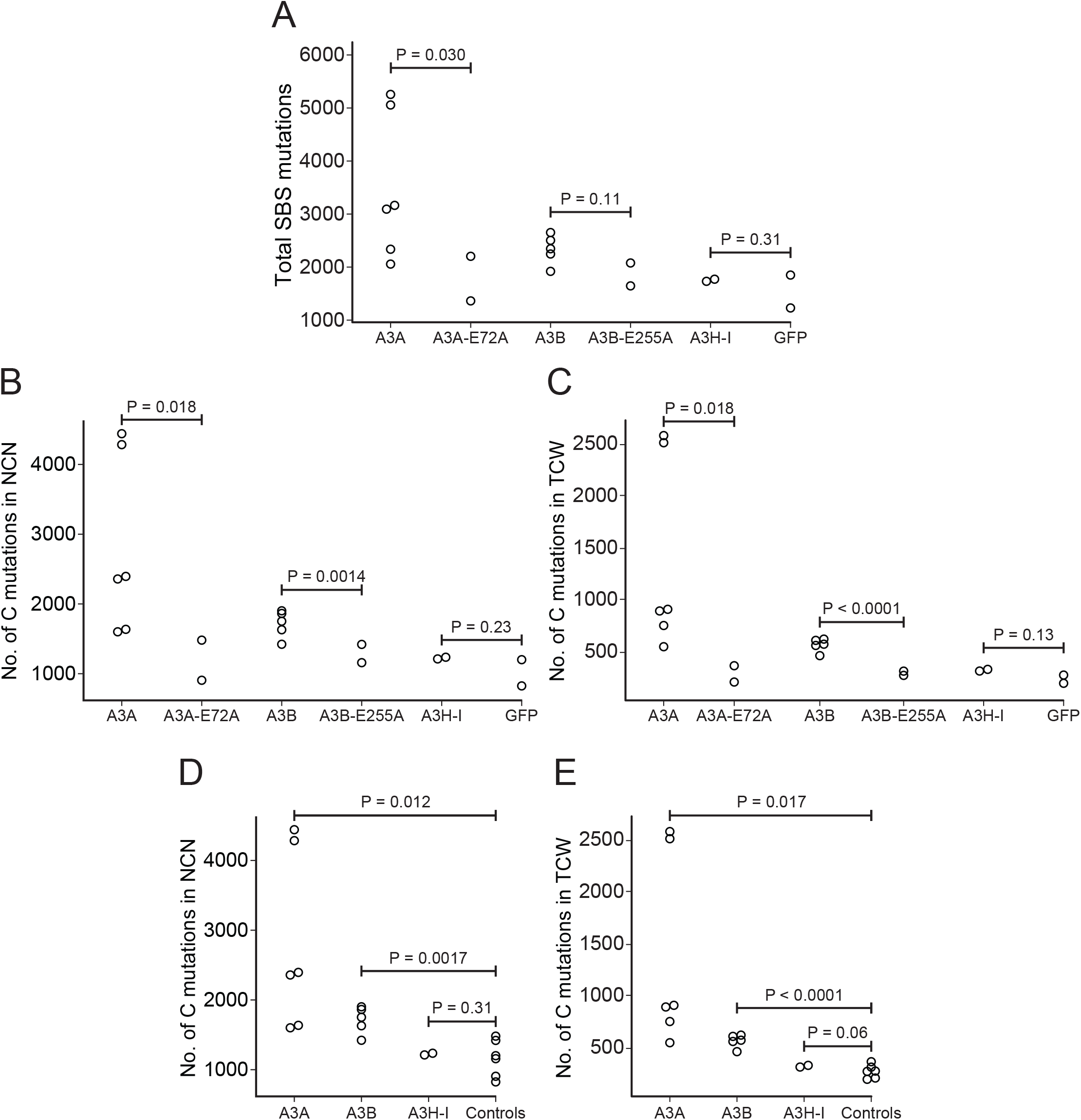
SBS mutation summary. (**A-C**) Dot plots showing total numbers of SBS mutations and cytosine mutations in NCN and TCW motifs, respectively, in WGS from individual Gan^R^ granddaughter clones (p-values using Welch’s t-test). (**D-E**) Dot plots showing total numbers of cytosine mutations in NCN and TCW motifs, respectively, in WGS from individual A3A and A3B expressing Gan^R^ granddaughter clones in comparison to all non-catalytic controls combined (p-values using Welch’s t-test).

**Figure 3 – Figure Supplement 5.**
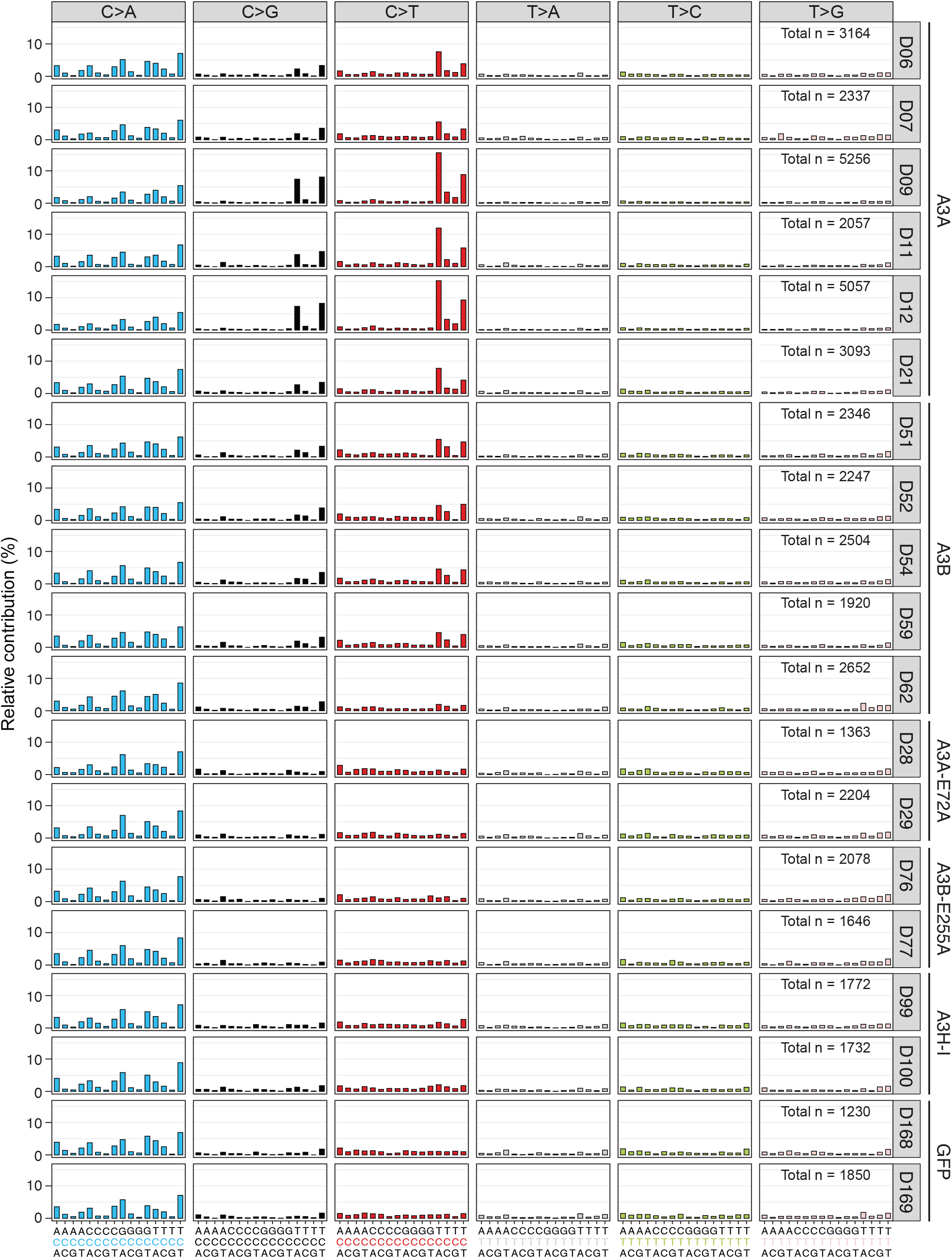
Single base substitution mutation profiles of individual ganciclovir-resistant clones by whole genome sequencing. Trinucleotide profiles of all SBS mutations in WGSs from the indicated Gan^R^ clones (conditions, clone names, and total SBS numbers are indicated to the right in each profile). Aggregate profiles for A3A, A3B, and corresponding catalytic mutant controls are shown in Figure 3.

**Figure 3 – Figure Supplement 6.**
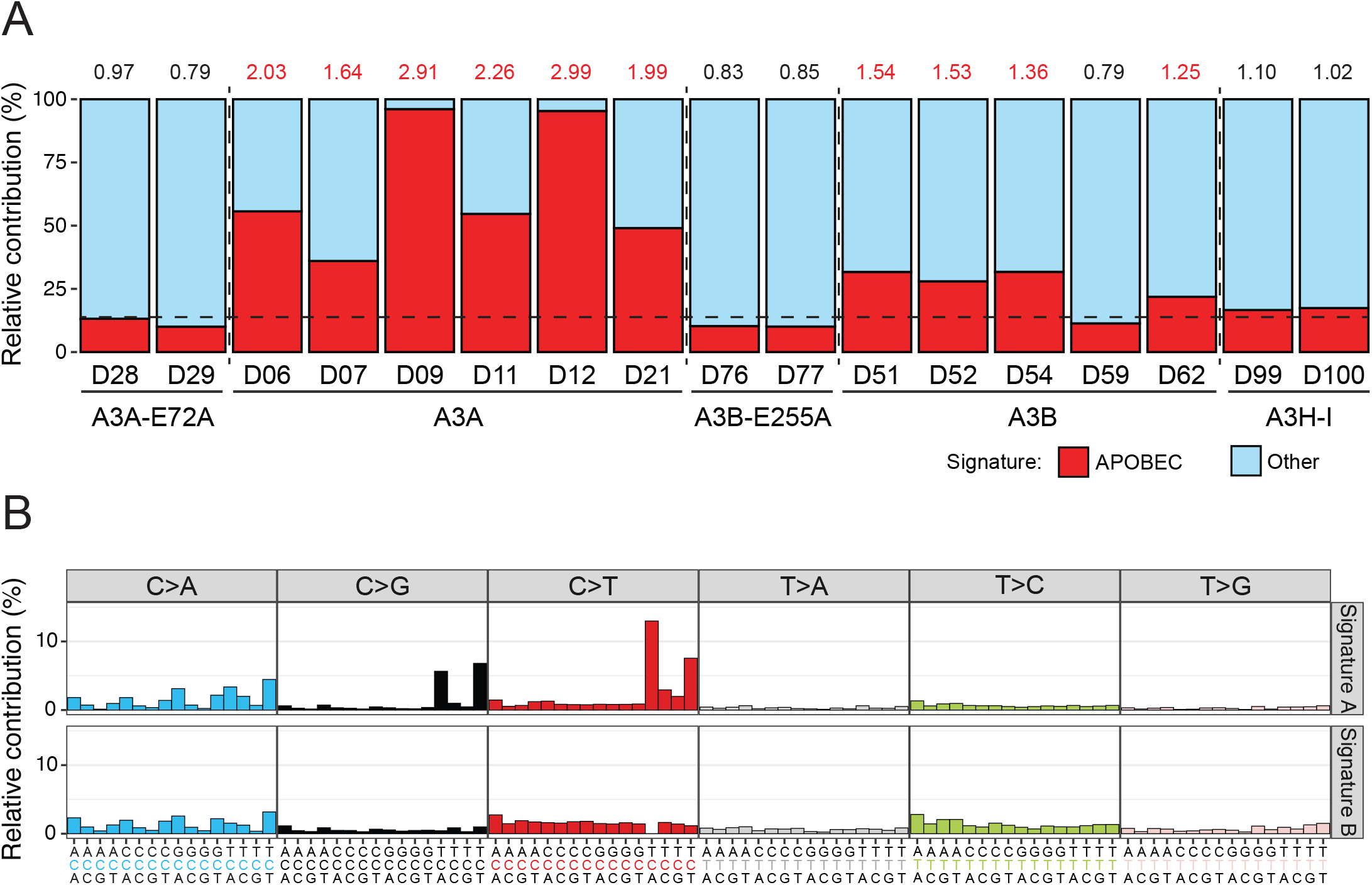
APOBEC3 mutation signature analysis using an alternative NMF-based workflow. (**A**) Mutation signature profiles extracted from Gan^R^ clone WGS using an NMF-based approach. The SBSs of each clone resolved into 2 signatures - an APOBEC3-like mutation signature shown in red and a background signature in blue. APOBEC3 mutation enrichment scores are shown above each bar, with significantly enriched values shown in red (Benjamini-Hochberg false discovery rate-corrected q-value < 0.05). The dashed line represents the average level of APOBEC3 signature mutations observed in the two eGFP control clones (*i.e*., background signal). (**B**) Trinucleotide mutation profiles of signatures A and B derived using NMF with the former exhibiting an APOBEC3 SBS signature (C-to-T and C-to-G in TCA and TCT motifs).

**Figure 3 – Figure Supplement 7.**
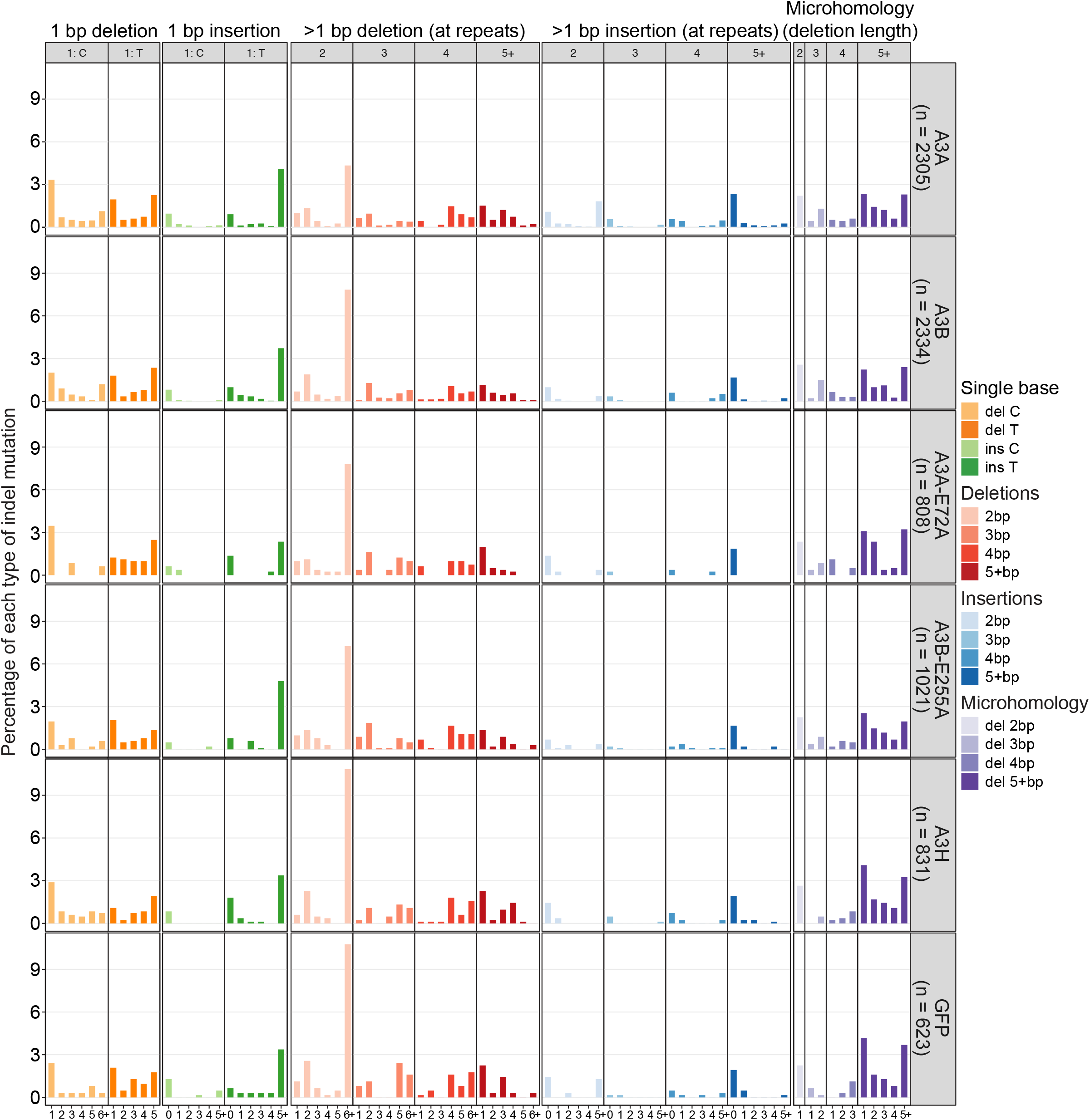
Indel landscape of sequenced granddaughter clones. Bar plots showing the percentage of each of the indicated indel types occurring in WGS from HAP1-TK-M9 granddaughter clones. Total numbers of single T deletions at homopolymers of 6 or more are too numerous to plot on the same axis and are therefore listed here (A3A, 39.5%; A3B, 44.2%; A3A-E72A, 40.6%; A3B-E255A, 40.0%; A3H, 31.8%; and GFP, 27.4%). The cosine similarity of the indel landscape across all conditions is over 0.96 indicating no significant differences.

**Figure 4 – Figure Supplement 1.**
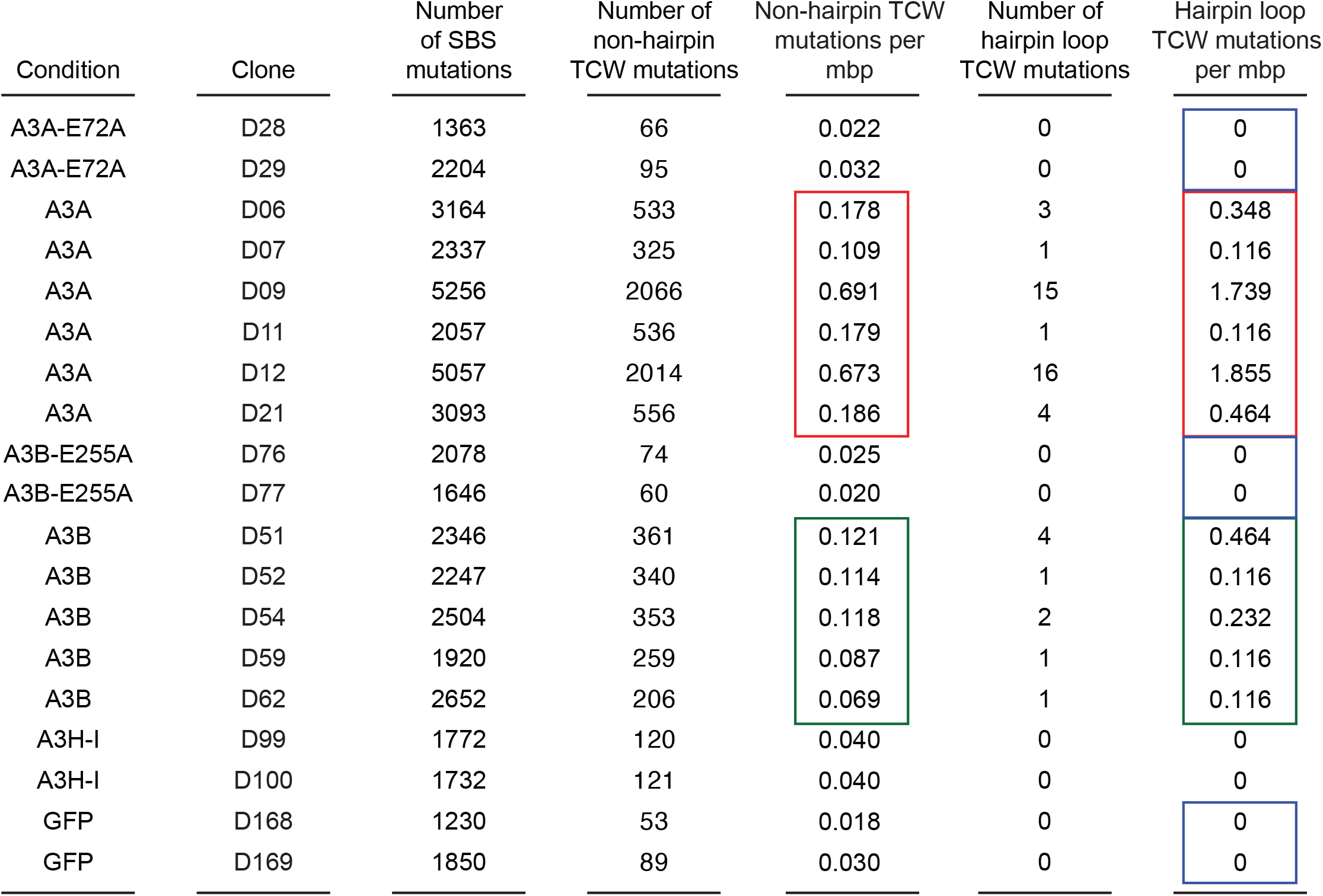
Hairpin mutation analysis. A list of the total number of SBS mutations in each HAP1-TK-M9 clone and numbers and frequencies mutations predicted to occur in non-hairpin regions or hairpin loop regions of the genome (see **Methods** for additional information). Columns 1 and 2: The A3 or control condition and clone number as listed in Figure 2 – Figure Supplement 1. Column 3: The total number of SBS mutations in each clone by WGS (human genome: 3000 mbp). Columns 4 and 5: The total number and frequency of non-hairpin APOBEC3 signature TCW mutations per clone (estimated non-hairpin genomic DNA: 2991.375 mbp). Columns 6 and 7: Total number and frequency of APOBEC3 signature TCW mutations in 3-11 nucleotide loop regions of predicted chromosomal DNA hairpin structures (estimated ssDNA loop region genomic DNA: 8.625 mbp). For A3A data (red boxes) and A3B data (green boxes), the non-hairpin versus hairpin APOBEC3 TCW mutation frequencies are not significantly different (P = 0.25 and P = 0.19, respectively, by Welch’s t-test). APOBEC3 TCW mutation frequencies are also not significantly different between the A3A and A3B data sets (red versus green boxes) for both non-hairpin regions (P = 0.087) or predicted ssDNA loop regions of hairpins (P = 0.149). However, in comparisons of A3A (red) and A3B (blue) APOBEC3 TCW mutation frequencies in ssDNA loop regions of hairpins and equivalent data sets from aggregate controls (blue boxes: the catalytic mutant of each protein and GFP), the A3A data set approaches statistical significance (P = 0.0655) and the A3B data are significantly different (P = 0.0367) by Welch’s t-test.

**Figure 4 – Figure Supplement 2.**
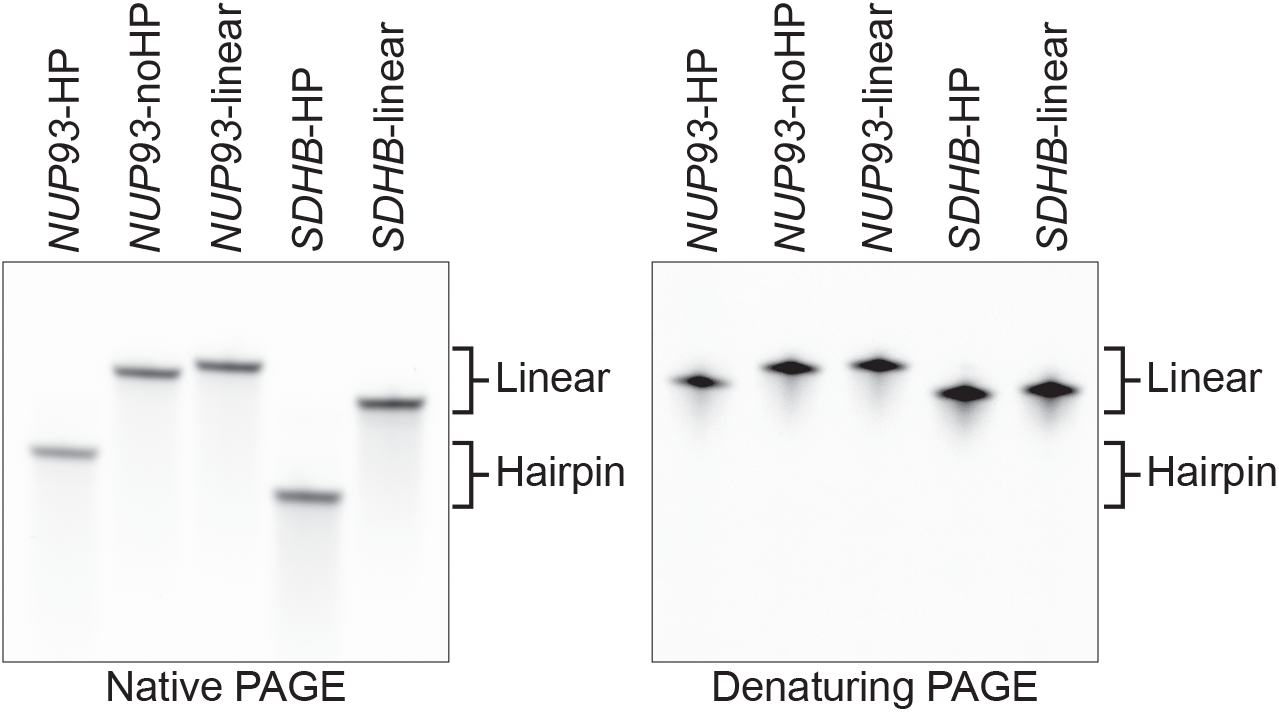
Validation of *NUP93* and *SDHB* DNA hairpins and linear structures. Native PAGE (left) and denaturing PAGE (right) analysis of the indicated oligonucleotide substrates. The hairpin substrates migrate faster under native conditions, and their mobility is similar to the linear derivatives under denaturing conditions. The only oligonucleotide not used in biochemical experiments in Figure 4 is the *NUP93*-noHP (no hairpin), which has half of the stem replaced by adenines (5’-6-carboxyfluorescein-GCAAGCTGTTCAAAAAAATGA) and is included here as an additional control.

**Supplementary Table S1. Pentanucleotide contexts of SBS mutations in HAP1-TK-M9 clones and ICGC breast tumors.**

This supplementary Microsoft Excel file reports the number of SBS mutations occurring in each sample in each pentanucleotide context. Layer 1 lists SBS mutations in Hap1-TK-M9 clones described here, and Layer 2 lists SBS mutations in ICGC primary breast cancers. The ICGC data are organized from left-most column to right-most column from highest APOBEC3 signature enrichment (lowest Benjamini-Hochberg q-value) to lowest APOBEC3 signature enrichment (highest Benjamini-Hochberg q-value). Columns reporting data from tumors with significant APOBEC3 mutation signature enrichments are shaded gray.

